# Lifestyle transitions in basidiomycetous fungi are reflected by tRNA composition and translation efficiency of metabolic genes

**DOI:** 10.1101/2023.05.09.539844

**Authors:** Marco Alexandre Guerreiro, Andrey Yurkov, Minou Nowrousian, Eva H Stukenbrock

## Abstract

Fungi are ubiquitous and inhabit every known terrestrial habitat. Pathogenic fungi are highly diverse, and recent years have seen an uprise in the emergence of new pathogens on crops, animals and humans. The order Trichosporonales (Tremellomycetes, Agaricomycotina, Basidiomycota) harbours saprobic and a few opportunistic human pathogenic species. These emerging pathogens cause superficial skin irritations, as well as invasive life-threatening infections. Yet, little is known about their evolution, ecology, virulence mechanisms and transition to pathogenic lifestyles. In this study we aimed to determine genomic signatures associated with lifestyle transitions, virulence and host/substrate specialization among 30 Trichosporonales species from a total of 41 genome sequences. We used comparative analyses of genome content, including gene functional categories, repetitive element content and tRNA composition among saprotrophic and reported opportunistic human pathogens. A genome-scale phylogenetic reconstruction revealed that even though the different genera are monophyletic, opportunistic pathogenic species are present in distantly-related clades. Statistical analyses showed that differences in genome structure among species did not correlate with predicted lifestyles. Intriguingly, we found that tRNA content varied widely across species (from 51 to 1455 manually curated tRNA genes). The expansion was independent from the phylogenetic structure. Opportunistic pathogenic species showed an overall increased efficiency in the translation of genes associated with host colonization (i.e. lipid metabolism), while exclusively saprotrophic species showed an increase translation efficiency for genes associated with a saprotrophic lifestyle (i.e. carbohydrate metabolism). This pattern was consistent among distantly-related saprotrophic and pathogenic *Cryptococcus* species (order Tremellales). In conclusion, our analyses link genomic information with ecology and fungal lifestyles across an entire order. We find evidence for an evolutionary scenario where distinct habitats select for an optimized translation of genes involved in successful proliferation in the respective habitat. We predict that lifestyles are not strictly defined by gene repertoires, but also by expression profiles in fungal pathogens.

## Introduction

Fungi are ubiquitous and inhabit every known terrestrial and aquatic habitat (Buzzini et al. 2017a, 2017b). Fungal individuals constantly interact with other organisms and with abiotic components of their habitat (Dighton 2016). These interactions are traditionally used to categorize their ecological role and lifestyle as saprotrophic, symbiotrophic or pathotrophic (Webster and Weber 2007). Depending on life stage and environmental conditions, many fungal species can undergo different lifestyles (Kuo et al. 2014). Due to the versatility of most fungal species, ecological roles are often circumstantial. As a consequence of these dynamic lifestyles, species classification into ecological groups is therefore usually challenging (Selosse et al. 2018; Kuo et al. 2014). Hence the drivers and mechanisms of lifestyle transitions in fungi are currently poorly understood.

Fungi are a worldwide fast-growing threat to human and plant health (Fisher et al. 2020; Fisher et al. 2012). Recent years have seen an uprise of newly emerging fungal pathogens on animals and plants (Fisher et al. 2012). Fungal pathogens are known to be highly diverse and to quickly adapt to host defences. Fungi often evolve very rapidly in response to new selective pressures in changing environments enabled by remarkable genomic and phenotypic plasticity. In clinical and agriculture scenarios, these unique properties grant fungi with the ability to infect new hosts, develop new infection mechanisms and virulence factors, and increased resistance to inhibitors (Jain et al. 2008; Giraud et al. 2010; Fisher et al. 2018).

Most human fungal pathogens are often found as part of natural environments or already associated with the host, as commensals or as pathogens (Brunke et al. 2016). The most common cases of fungal infection in humans are caused by members of *Aspergillus*, *Blastomyces*, *Coccidioides*, *Cryptococcus*, *Candida*, *Histoplasma*, *Malassezia*, *Paracoccidioides*, *Penicillium*, *Pneumocystis*, *Rhizopus* and *Trichosporon* (Guého et al. 1994; Brown et al. 2012; Schwartz 2004; Almeida Júnior and Hennequin 2016; Fisher et al. 2020). While some fungal species are highly adapted to human hosts, e.g. *Candida*, *Malassezia* or *Trichophyton*, many species have an environmental origin and are usually considered as opportunistic pathogens such as species of *Cryptococcus*, *Trichosporon*, *Histoplasma*, *Blastomyces* or *Aspergillus*.

A key feature that defines pathogens is the ability to survive and disseminate in the host physiology and body temperature (Casadevall 2012). Accordingly, opportunistic human pathogens are often equipped with evolutionary adaptions that enable them to colonize and survive in both natural environments and human hosts (Hube 2009; Brunke et al. 2016). For example, the opportunistic human pathogens *Cryptococcus deneoformans* and *Cryptococcus gattii* (Basidiomycota), responsible for systemic infections, show ability to grow up to 40 °C, while the saprobic sister species *Cryptococcus amylolentus* and *Cryptococcus floricola* only grow at temperatures up to 30 °C (Fonseca et al. 2011; Kwon-Chung 2011; Passer et al. 2019).

Fungal pathogens must interact with the host defence and avoid immune systems (Voelz and May 2010; Grice and Dawson 2017). Through evolution, selection and adaption, pathogenic and opportunistic fungi have developed virulence factors that grant them the ability to colonize different organisms and disseminate while in association with the host, often causing infectious diseases. Virulence factors are usually related to morphological plasticity and dimorphism, biofilm production, thermotolerance, metabolism, enzyme production and secretion (Hogan et al. 1996; Jain et al. 2008; Ramage et al. 2009; Alspaugh 2015; Boyce and Andrianopoulos 2015; Bielska and May 2016; Gauthier 2017; Muñoz et al. 2018). Some of these virulence factors are specific to lifestyles and hosts, and can in some cases be assessed based on genomic or physiological data (Xu et al. 2007).

Some well-defined lifestyles and fungal groups have been reported to be associated with specific genomic features, including gene content. These include the loss of genes encoding plant cell-wall degrading CAZymes in mycorrhizal fungi (Kohler et al. 2015; Peter et al. 2016; Miyauchi et al. 2020), expansion of CAZyme repertoire in saprotrophic and wood-degrading fungi (Rytioja et al. 2014), or secretion of host-specific effector proteins in fungal phytopathogens (Dodds and Rathjen 2010; Haueisen and Stukenbrock 2016). All members of the genus *Malassezia*, which comprises anamorphic species adapted to the human skin, lack genes encoding for fatty acid synthases; however, this is counterbalanced by an expansion of genes encoding for secreted lipases and phospholipases, which are used to acquire lipids from the host (Xu et al. 2007).

Several studies have documented that changes in gene expression patterns are core components of adaptation to new environments (Yona et al. 2013). This can be demonstrated by comparative analyses of transcriptomes. Additionally, it can be reflected in the tRNA pool that rapidly evolve to meet new translational requirements by mutating more abundant and redundant anticodons. This level of adaptation involves changes in the relative numbers of the different tRNA molecules encoded by the genome (Yona et al. 2013). The tRNA-mediated processes have been shown to provide valuable insights into the evolutionary mechanisms of fungal adaptation (Badet et al. 2017), in addition to traditional studies of gene expression patterns (Yona et al. 2013).

tRNA genes are classified into families according to their anticodon. Each tRNA gene family may be present as a single gene or as several copies scattered throughout the genome. In the latter case, multiple copies are considered to be functionally redundant due to the same anticodon sequence. Studies have empirically demonstrated that in fungi and bacteria, the tRNA copy number in the genome correlates with the abundance of each tRNA family in the cell (Dong et al. 1996; Percudani et al. 1997; Kanaya et al. 1999; Tuller et al. 2010). Therefore, gene copy number in the genome can be used as an estimation for the available tRNA pool in the cell.

Protein synthesis is in part regulated by the availability of tRNAs that match the codons being translated. This regulation is possible due to codon usage biases, where synonymous codons are used in unequal frequencies throughout the genome (Parvathy et al. 2022). This fine tune between the available tRNAs and the codon usage bias allow the regulation of the protein expression of individual genes. Highly expressed genes have an optimal codon composition that correspond to abundant tRNAs, which promote a faster and more efficient translation (Sun et al. 2016). Synonymous mutations in coding-regions that alter the codon usage bias can therefore be selected during evolution based on fitness advantage (Supek 2016; Frumkin et al. 2018). Such evolutionary forces ensure an optimal translation efficiency of essential genes. tRNA gene redundancy and genome size have been determined as key factors for translational selection, indicating that optimal translational effiency requires a fine tuning between these two factors (dos Reis et al. 2004). Codon frequencies and tRNA copy number may therefore co-evolve to maintain an optimum balance for optimal protein synthesis (Higgs and Ran 2008; Gingold et al. 2012). Codon frequencies and tRNA copy number can therefore be used to estimate how efficiently genes may be translated (dos Reis et al. 2004) and thus estimate protein synthesis rates, cell response to external factors and ultimately, evolutionary adaptations to new environments. Codon optimization has been shown to underlie genome adaptation in fungal pathogens and to be linked to their ability to colonize multiple hosts (Badet et al. 2017). So far, little is known about fungal lifestyle dynamics and the evolutionary mechanisms granting rapid transitions between lifestyles in some species.

The basidiomycetous order Trichosporonales comprises many species widely distributed in nature. This order includes various species that have clinical, agricultural, and biotechnological importance (Sugita 2011). Currently, there are no sexual stages reported for these species, but the genes required to undergo sexual mating have been identified in all studied genomes (Sun et al. 2019). Trichosporonales are emerging pathogens, responsible for superficial skin irritations, but also cause invasive life-threatening systemic infections. However, little is known about their evolution, ecology, virulence mechanisms and the transitions to pathogenic lifestyles. The genus *Trichosporon* (Trichosporonales) contains many known pathogens of animals and humans (Sugita 2011). Superficial and invasive infections by *Trichosporon* species are emerging life-threatening conditions (Guého et al. 1994; Schwartz 2004; Almeida Júnior and Hennequin 2016). Invasive infections by *Trichosporon* are frequently fatal for immunocompromised patients, with an estimated 77% mortality rate (Girmenia et al. 2005). Despite its epidemiological significance, only a few pathogenicity traits have been reported and addressed for this genus, such as biofilm production, broad antimycotic resistance and growth at high temperatures (Chakrabarti et al. 2002; Di Bonaventura et al. 2006; Pagano et al. 2006).

Currently, adaptive traits related to pathogenicity have not been identified in this genus or related genera. Since closely related species exhibit different lifestyles alternating between pathogenic and saprotrophic, an intriguing question is moreover how pathogenicity related traits evolve.

In this study, we aimed to determine genomic signatures associated with lifestyle, virulence and host/substrate specialization in fungi. We applied a set of closely related Trichosporonales species as model system allowing us to address genomic changes associated with recent life-style transitions. For that, we characterized and compared the genome structure, functionality and translational adaptation among saprotrophic and reported opportunistic human pathogens. Due to the diverse and polyphyletic ecologies in this order, we focused on identifying signatures granting an opportunistic pathogenic lifestyle. Due to their saprotrophic and dermatophytic occurrences, we focused on their carbon and lipid transport and metabolic pathways to further characterize lifestyles.

We hypothesized that (i) genomes of opportunistic pathogens will have a higher diversity and copy number of genes associated with lipid transport and metabolism when compared to genomes of saprotrophic species, but (ii) similar content of genes related to carbohydrate transport and metabolism, including CAZymes. Additionally, we hypothesized that (iii) lifestyle adaptations are imprinted in the translation machinery, namely in tRNA pools and codon optimization in pathways relevant to the lifestyle. Due to the ecological versatility of these species, (iv) some plasticity is expected in terms of gene regulation and translation, as well (v) independent and convergent evolutionary events among species with similar ecological strategies. Using information from 41 genomes of Trichosporonales species, we provide evidence that opportunistic fungal pathogens are not strictly defined by gene repertoires, but rather by their ability to readily evolve and adapt to new environments through adaptive translation.

## Materials and Methods

### Genomic Data Acquisition

Assembled genomic data of 50 fungi were downloaded from the NCBI database, US DOE Joint Genome Institute’s MycoCosm database (https://mycocosm.jgi.doe.gov) or ATCC Genome Portal (https://genomes.atcc.org/genomes). A total of 5 hybrid or diploid genomes were excluded from further analyses based on literature (Takashima et al. 2018) and duplicated BUSCO content (Fig. S1). Strains with a predicted proteome completeness higher than 85% and duplicated BUSCOs lower than 5% were selected for further analyses. The final dataset comprised a total of 41 haploid genomes, representing 30 species. Additionally, 4 *Cryptococcus* species were considered as outgroups.

Species were considered as opportunistic human pathogens if they were 1) previously reported in human clinical settings and 2) the identify confirmed with molecular markers (Table S1). Species without any clinical report were considered as saprotrophic. Due to the wide environmental occurrence of these fungi (e.g. plant-associated, water samples, insects, soil, dairy), the species lifestyle was considered for the analyses rather than the strain isolation substrate.

### Identification of species boundaries

Average nucleotide identities were calculated by performing pairwise whole-genome alignments with MUMmer v3.23 (Kurtz et al. 2004) and a delta filtering step with ’-1’ and ’-l 100’ parameters. Species boundaries were defined based on barcode and genome average nucleotide similarity. To reduce biases while comparing genomic features among lifestyles, one representative for each species with the most contiguous genome assembly was selected for statistical analyses.

To verify the genome species identification and intra-specific diversity, the ribosomal RNA regions, corresponding to the molecular barcodes ITS and LSU were retrieved from the genome assemblies. These barcode regions were subsequently compared to type-strains and among the isolates in study via BLAST and manually curated alignments. When absent in the genome assembly, the barcode regions were retrieved and assembled based on the respective raw reads. Raw reads were downloaded (Table S1) and quality-trimmed with Trimmomatic v0.39 (Bolger et al. 2014). The quality-filtered reads were mapped against a set of reference barcode sequences with BBmap tools v38.76 (Bushnell 2017), using default settings, which allow a minimum alignment identity of 0.76 and 32 maximum mismatches. Mapped reads were assembled with SPAdes v3.15.0 (Bankevich et al. 2012). These resulting assemblies were then compared to the abovementioned barcode dataset, containing type-strains and other strains analysed in this study (Table S2).

### Coding gene prediction

To avoid gene prediction biases, all assembled genomes were processed using the funannotate pipeline v1.8.10 (Palmer and Stajich 2020), with the same settings. The pipeline comprised size filtering, cleaning, sorting and softmasking steps with default parameters, followed by gene prediction using the “cryptococcus” as *augustus_species* and *busco_seed_species* and “basidiomycota” as *busco_db*. Genome completeness was predicted using BUSCO v5.3.2 (Manni et al. 2021) in comparison to the Basidiomycota lineage database (basidiomycota_odb10). For gene comparisons, only coding sequences (CDS) without a predicted stop codon in the inner sequence were considered for further analyses.

### Phylogenetic reconstruction

Orthologous protein sequences among the studied Trichosporonales were predicted by using OrthoFinder v2.5.4 (Emms and Kelly 2019). A set of 1438 single-copy orthologous genes, present in all species, were obtained and were subsequently individually aligned with MAFFT v7.505 (Katoh and Toh 2008). The individual gene alignments were concatenated and used to calculate a maximum likelihood phylogenetic tree with RAxML v.8.2.12 (Stamatakis 2014), using 500 bootstrap replicates, the PROTGAMMAWAG model, 123 as seed number for the parsimony inferences and a random seed of 321.

### tRNA gene prediction and analyses of sequence variation

tRNA genes were predicted with tRNAscan-SE 2.0.9 (Chan et al. 2021) with defaults parameters for eukaryotic organisms. To assure high-quality of gene predictions, all possible pseudogenes and tRNA genes with a covariance score below 30 (Chan et al. 2021) were excluded from further analyses. Additionally, tRNA genes were manually curated based on their predicted structure. tRNA gene copy number was considered as a proxy for tRNA abundance.

To estimate sequence variation, tRNA gene sequences were obtained for each genome and aligned for each anticodon type by using MAFFT v6.864b (Katoh and Toh 2008). Only genes with two or more copies were considered for intragenomic sequence variation analyses. To estimate intragenomic sequence variation for each tRNA gene family (i.e. anticodon type), the genetic distance was predicted based on sequence alignments using the Kimura-2-parameter distance model (Kimura 1980) in the *ape* R package v5.6-2 (Paradis and Schliep 2019). Outliers with a distance above 10 were excluded, which likely resulted from poor alignments.

### Prediction of additional genomic features

To predict CAZymes, proteomes were scanned against dbCAN2 v10.0 database (Zhang et al. 2018) using HMMER v3.3 (Eddy 2011). Matches with an e-value lower than 1e-15 and a coverage higher than 0.35 were used for further analyses. Secreted proteins were predicted from each proteome with SignalP 6.0g (Teufel et al. 2022). Transposable elements were predicted by analysing each genome assembly with REPET3 pipeline (Quesneville et al. 2005; Flutre et al. 2011) and according to Lorrain et al. (2021). Repetitive elements were determined by funannotate during soft masking with tantan (Frith 2011).

### Codon optimization and tRNA adaptation index estimation

To infer translational selection, the tRNA adaptation index (tAI) and codon optimization index (S) were calculated with the *tAI* v0.2 R package (dos Reis et al. 2004). The tRNA adaptation index (tAI) measures the co-evolution between a given coding sequence and the respective genomic tRNA pool (dos Reis et al. 2003; dos Reis et al. 2004). The codon optimization index (S) estimates the intensity of translational selection based on the correlation between the tAI and the effective number of codons (corrected for GC content at third codon positions) (dos Reis et al. 2003; dos Reis et al. 2004). The S index therefore reflects the intensity of translational selection on gene sets and respective translation efficiency.

### Phylogenetic signal estimation

The strength and direction of trait evolution was estimated based on Blomberg’s K (Blomberg et al. 2003), using phylogenetic distance as the only predictor of trait similarity among species. Blomberg’s K infers stabilizing selection (K>1) and neutral (K=1), or convergent (K<1) evolution of a trait, relative to the expected variation under the Brownian Motion model of evolution (Blomberg et al. 2003). This parameter was calculated using the *phytools* v1.0-3 R package (Revell 2012) and using 100 permutations. Only K values with a *P* < 0.05 were considered as significant. A previously calculated maximum likelihood phylogeny with branch length was used for the calculation (see above).

### Statistical analyses

Statistical analyses were performed in R v4.2.0 (R Core Team 2022) and RStudio 2022.02.2 (RStudio Team 2022). Data was visualized with *ggplot2* (Wickham 2016). For correlative analyses, only one

strain of each species was used. Wilcoxon tests were performed with *ggsignif* (Ahlmann-Eltze and Patil 2021) and *broom* (Robinson et al. 2022) R packages. Linear models, R-values and respective significance *P*-values were calculated with *ggpmisc* R package (Aphalo 2022).

### Data availability

The functional annotation pipeline, R scripts and additional scripts used to create the results presented in this manuscript can be found at https://github.com/maguerreiro/Trichosporonales

## Results

### Distinct lifestyles among closely related species of Trichosporonales

To determine the evolutionary relationship among members of the Trichosporonales, we inferred the species phylogeny based on 1438 single-copy orthologous protein sequences present among all strains. Furthermore, we compared the resolution for species discrimination of the phylogenomic approach with the barcoding genomic ribosomal RNA regions comprising the ITS region (containing the ITS1, 5.8S, ITS2) and the LSU region (containing the D1/D2 region of the 26S ribosomal large subunit). This approach was further used to confirm the species identity of the genomic data not belonging to type-strains.

The phylogenomic tree showed an overall 100% bootstrap support for the backbone, except for the clade containing the Pascua and Prillingera genera, which were supported by 98% (Fig. 1). The topology of the tree indicates that the different genera are monophyletic. However, the Apiotrichum genus is divided into two distinct lineages, with *Apiotrichum porosum* and *Apiotrichum gamsii* being separated from the remaining 9 *Apiotrichum* species. A phylogenetic principal coordinate (pPCo) analysis, based on pairwise phylogenetic distances (i.e. branch length) between species, reflected the topology of the phylogenetic tree (Fig. 2). This non-hierarchical analysis supported the separation of the two Apiotrichum lineages. Furthermore, the phylogenetic distance between the two Apiotrichum clades (pairwise phylogenetic distances = 0.611 – 0.691) was similar to the distances between different genera (Table S3), e.g. *Apiotrichum porosum* and *Prillingera fragicola* (0.687) or *Haglerozyma chiarellii* and *Vanrija humicola* (0.675), suggesting that these two Apiotrichum lineages might belong to two different genera (Fig. 2). Additionally, pPCo analysis revealed the relatedness among species in the Trichosporon and Cutaneotrichosporon genera, which overlap in terms of phylogenetic clustering and which share common lifestyles, suggesting that these might be part of a species complex (Fig. 2). Furthermore, the genus Cutaneotrichosporon revealed a high phylogenetic distance among species (up to 0.805), which could indicate, at least, 3 different lineages (Fig. 2, Table S3).

**Figure 1.**
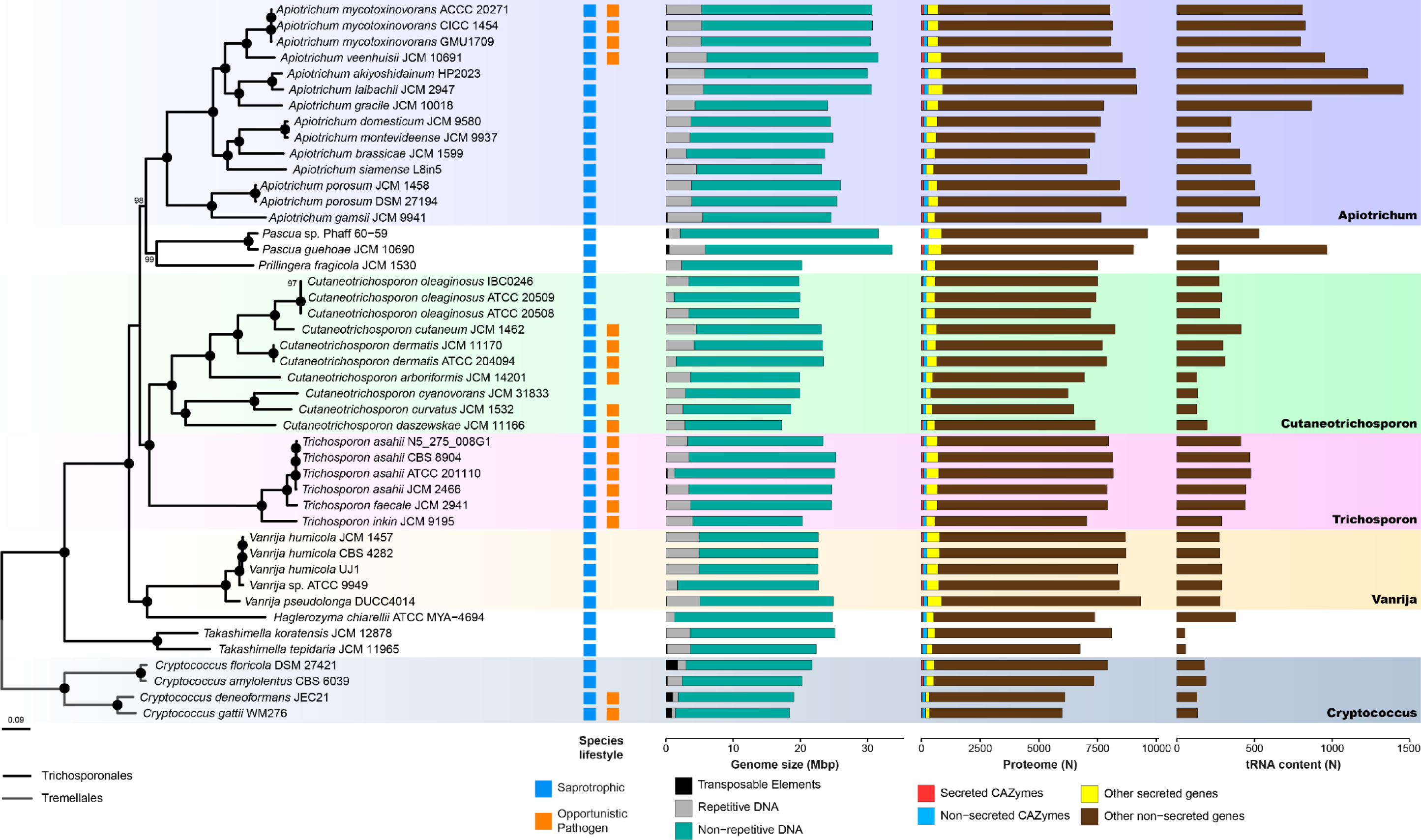
Overview of the haploid Trichosporonales species analysed and their respective genomic features. A maximum likelihood genome-scale phylogeny of the Trichosporonales and Tremellales was calculated based on the protein sequence of 1438 single-copy orthologous genes. Lifestyles of the species used in this study are colour-coded for saprotrophic (blue) and opportunistic pathogenic (orange) species. The genome size for each genome assembly is indicated, with the respective fractions assigned as transposable elements (black), repetitive sequences (grey) and non-repetitive DNA (green). The predicted proteome for each strain is indicated and divided into secreted CAZymes (red), non-secreted CAZymes (blue), additional secreted genes (yellow) and additional non-secreted genes (brown). In the last column, the total number of detected tRNA genes is depicted.

**Figure 2.**
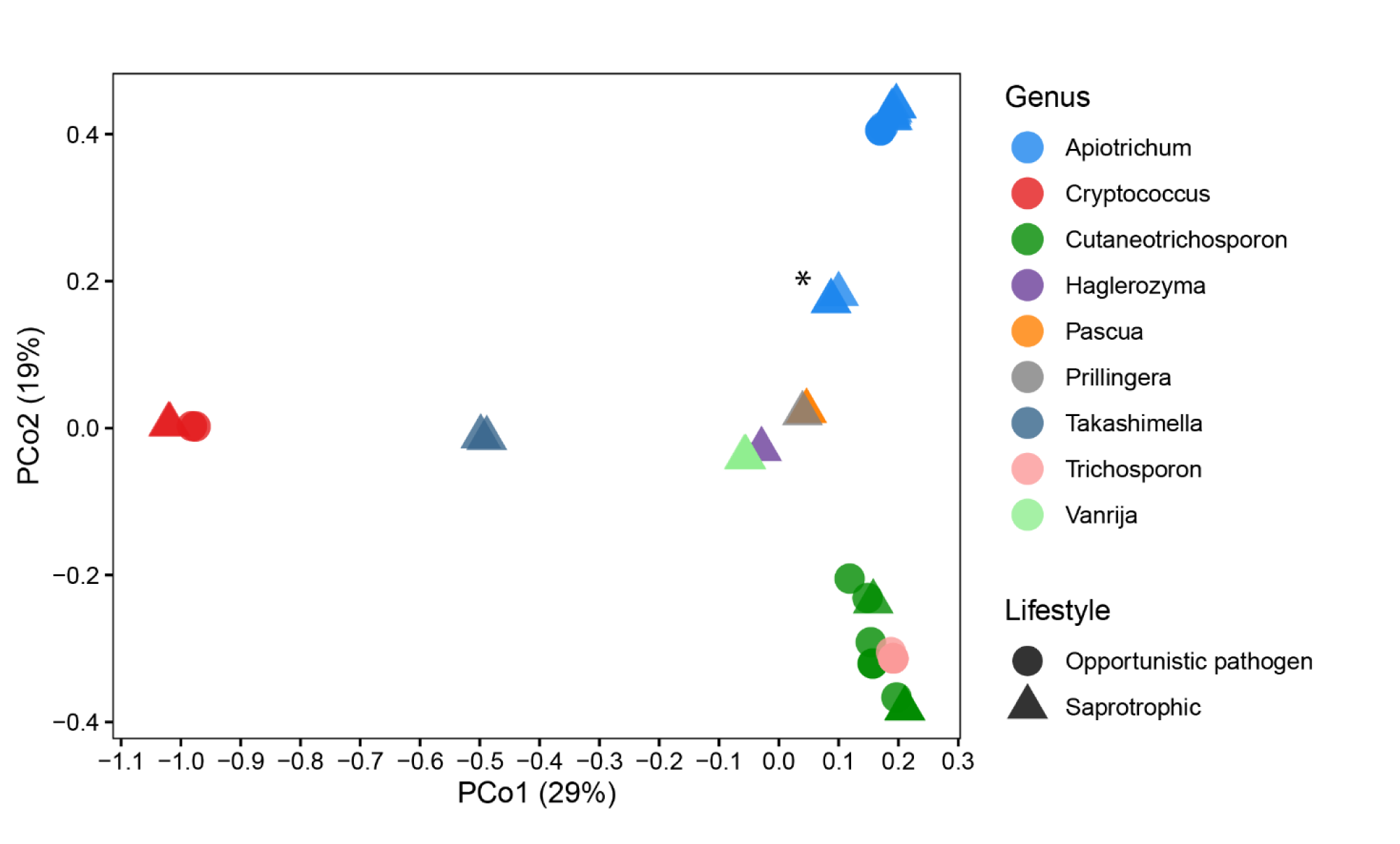
PCoA of the genomic pairwise phylogenetic distances between individuals. Genera are colour-coded and lifestyles are coded by symbols. The asterisk (*) indicates the position of *Apiotrichum gamsii* and *Apiotrichum porosum* genomes.

The phylogenetic analyses additionally revealed that opportunistic pathogenic species are located mainly in the Trichosporon and Cutaneotrichosporon clades, with two species belonging to Apiotrichum (Fig. 1). The dispersion of pathogenic species across the phylogeny implies that the emergence and loss of pathogenicity have occurred several times in the Trichosporonales order.

In addition, the phylogenetic analyses also revealed unexpected divergence between individuals of *Pascua guehoae* and *Vanrija humicola*. One genome assigned as *Pascua guehoae* (Phaff 60-59) and one of *Vanrija humicola* (ATCC 9949), showed high genome dissimilarity to related genomes. This could indicate a high intraspecific variation or a misidentification of the species in these strains. To clarify this, we compared the averaged genome similarity with the similarity of molecular barcodes, which are conserved within species and used to identify and delimit species.

Phaff 60-59 showed an averaged genome nucleotide similarity of 87.8% to the genome of another isolate assigned to the same species, *Pascua guehoae* (JCM 10690^T^), the closest phylogenetic related species and the species type-strain (Table S3). In addition, Vanrija ATCC 9949 showed a genomic similarity of 90.8% to the several *Vanrija humicola* isolates (CBS 571^T^, CBS 4282 and UJ1). For comparison, both *Apiotrichum porosum* strains showed a 97.2% nucleotide similarity (Table S3), while pairwise comparisons between different species on genome-scale yielded a maximum of 92.11% similarity between *Apiotrichum domesticum* and *Apiotrichum montevideense* (Table S3). To further assess the taxonomic status of these strains, we retrieved and compared the similarity among the ribosomal barcoding regions ITS and LSU.

In the retrieved barcode regions, ITS and LSU, Phaff 60-59 differed by 5 nucleotides (0.5%) from JCM 10690^T^, while ATCC 9949 differed by 8 nucleotides (0.8%) from CBS 571^T^ (Table S4). The comparison between both *Apiotrichum porosum* strains showed that the barcodes were identical (100% similarity), while *Apiotrichum domesticum* and *Apiotrichum montevideense* differed by 3 nucleotides (0.3%). This suggests that both Phaff 60-59 and Vanrija ATCC 9949 strains have a dissimilarity to related strains at a higher level than that observed between different species.

In summary, we observed that a dissimilarity of 0.8% on the barcodes between ATCC 9949 and *Vanrija humicola* CBS 571^T^ corresponded to an average nucleotide dissimilarity of 9.2% on a genome-wide scale (Table S5). Likewise, a 0.5% difference in these barcodes between Phaff 60-59 and *Pascua guehoae* JCM 10690^T^ corresponded to a 12.2% average nucleotide dissimilarity between both genomes. Strains with 100% barcode identity, differed up to 2.8% across the whole genome (Table S3 and Table S4). These results suggest that the isolates *Pascua* sp. Phaff 60-59 and *Vanrija* sp. ATCC 9949 might represent undescribed species. Furthermore, our results highlight the importance and advantage of comparing barcode and genome-wide similarity among strains to verify taxonomic assignments and help defining species boundaries from a phylogenetic perspective. Comparison of multiple strains per species with identical barcodes (100% identity) indicate that species might be delimited at 97% genome-wide averaged nucleotide similarity.

### Saprotrophic and opportunistic pathogens in the Trichosporonales are indistinguishable based on genomic features

In order to identify genomic signatures associated with lifestyle and identify signatures of convergent lifestyle-induced evolution, we characterized and compared the genome structure and composition between opportunistic pathogens and saprotrophic species. To categorize species as saprotrophic or opportunistic pathogens, we used information from previously published clinical reports (Table S1). We classified species as opportunistic human pathogens when these were reported from human clinical settings and as saprotrophic species when no clinical occurrence was reported. To avoid statistical biases, we selected one representative strain per species with the most contiguous genome assembly (Table S1).

Based on a high content of duplicated BUSCOs, five species were predicted to be hybrids or diploids, and therefore removed from further analyses (Fig. S1). Genome completeness, based on complete BUSCOs, of the analysed assemblies ranged from 85.2% (*Pascua* sp. Phaff 60-59) to 98.6% (*Apiotrichum laibachii* JCM 2947). The haploid genome assemblies among the Trichosporonales species were estimated to range from 17.22 Mb (*Cutaneotrichosporon daszewskae* JCM 11166) to 33.70 Mb (*Pascua guehoae* JCM 10690). Correlating genome size to the content of transposable elements (TEs) in the genome, we found a significant correlation (R² = 0.58, *P* = 0.00015), suggesting that TEs composition is a determinant of genome expansion. On the other hand, there was no correlation of genome size with simple repeat content (*P* > 0.05) (Fig. S2).

We next asked if the predicted fungal lifestyle was reflected in genome structures or the content of genomic features (i.e. genome size, total number of genes and secreted genes, GC content, total number of CAZymes and secreted CAZymes, total number of tRNA genes, repeat and TE content). These features vary considerably between species, however, based on Wilcoxon, PERMANOVA and ANOSIM tests, we found no significant differences in genome structure and the content of genomic features among saprotrophic and opportunistic pathogens members of the Trichosporonales (Fig. 3). These results support our initial hypothesis (ii), that saprotrophic and opportunistic pathogenic species harbour similar CAZyme content, due to their occurrence in natural environments. Intriguingly, we also find that the number of genes encoding for secreted proteins is comparable between pathogenic and saprotrophic species. Often secreted proteins are considered as putative virulence factors. The similar composition of secreted proteins suggests that saprotrophic species may have a virulent potential or that these proteins may play other roles beyond virulence.

**Figure 3.**
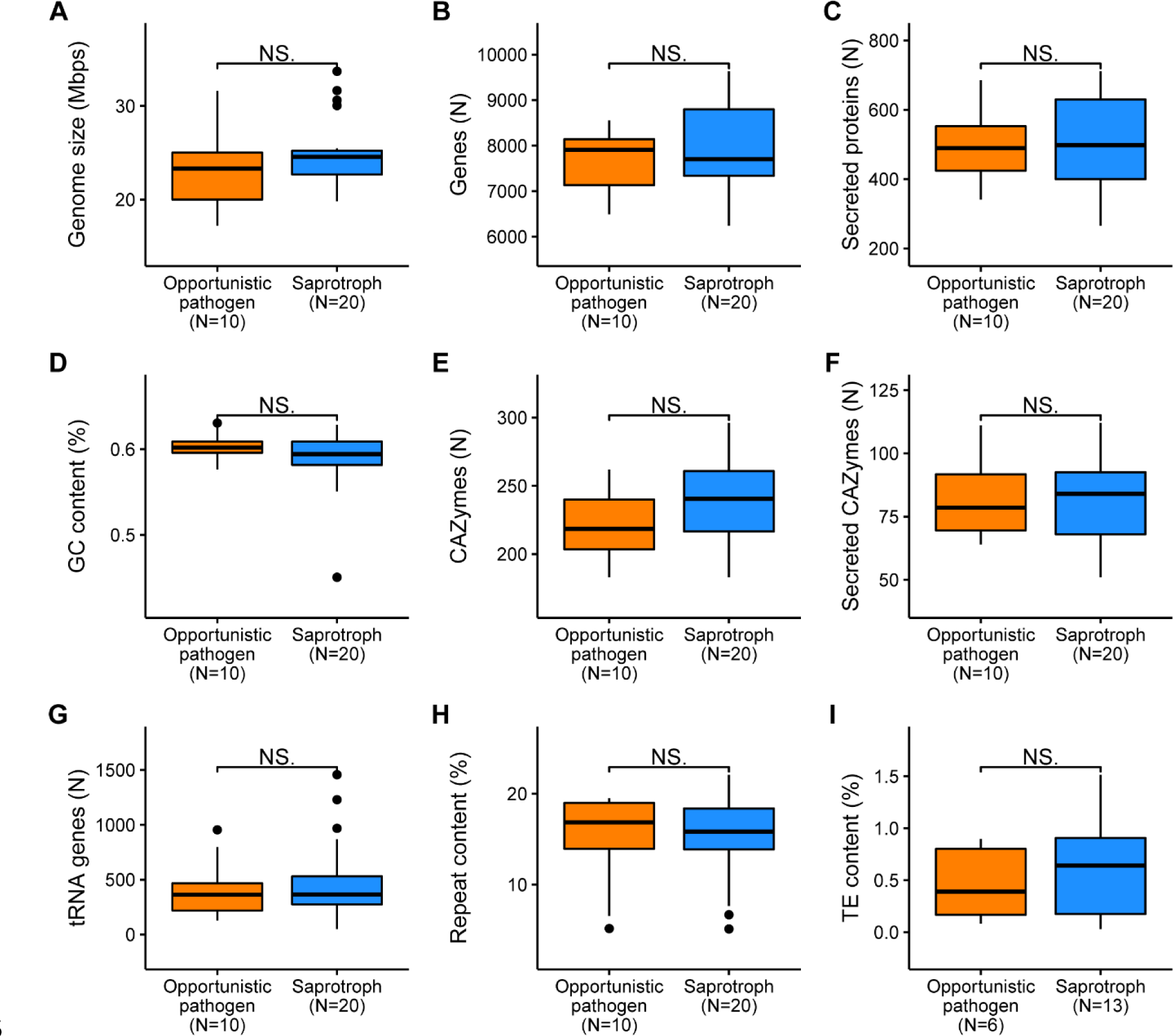
Comparison of genomic features between opportunistic pathogenic and saprotrophic lifestyles among species in the Trichosporonales order. No significant statistical differences (“NS.” *P*-value ≥ 0.05) were observed based on Wilcoxon signed-rank test. To avoid overestimation, only one genome per species was considered and the total number of included genomes is indicated.

We note that a few patterns were observed when the same analyses were performed on single genera of Trichosporonales and Tremellales, containing both lifestyles. However, the low sample size did not allow for statistical inferences. Saprotrophic *Cryptococcus* species generally encoded more secreted proteins and CAZymes than opportunistic pathogenic species (Fig. S3). These differences were however not consistent among *Apiotrichum* and *Cutaneotrichosporon* species (Fig. S3). This indicates that the evolutionary history of these genomic features might be different among these genera.

### tRNA composition and expansion among Trichosporonales are independent from the phylogenetic structure

Due to the high variation of tRNA content among species (Fig. 1), we further investigated the distribution of tRNA genes and their expansion along the species phylogeny.

Comparative analyses revealed that the total number of tRNA genes (tRNAome) varied widely across the 41 analysed Trichosporonales genomes (Fig. 1 and Table S6). The mean tRNAome comprises 444 genes, with a minimum of 51 and a maximum of 1455 tRNA genes (Fig. S4). Hereby, *Apiotrichum laibachii* and *A. akiyoshidainum* possessed unusual large tRNAomes with 1455 and 1227 genes, respectively, while *Takashimella koratensis* and *Ta. tepidaria* possessed comparatively much smaller tRNAomes with 51 and 58 genes, respectively (Table S6). These analyses revealed an uneven and wide distribution of tRNA gene content along the phylogeny.

We again asked if fungal lifestyle was reflected in the content of tRNA genes. However, we find that the tRNAome size was indistinguishable between opportunistic pathogenic and saprotrophic lifestyles (Fig. 3G). The tRNAome was strongly correlated with the genome size (Fig. S5) among all Trichosporonales (R² = 0.60, *P* = 5.3e−07). This correlation was stronger among opportunistic human pathogenic species (R² = 0.93, *P* = 7.0e-06) compared to saprotrophs (R² = 0.52, *P* = 3.2e-04), which may suggest a stronger selection and finer tune between tRNA genes content and genome size among opportunistic human pathogenic species.

The number of the detected unique anticodon types (i.e. tRNA gene families) ranged from 39 in *A. porosum* (JCM 1458) and *Ta. koratensis* to 46 in *A. mycotoxinovorans* and *Haglerozyma chiarellii* (from the universal 61 possible anticodons) (Fig. S6 and Table S6). Nevertheless, we were able to detect tRNAs decoding all 20 universal amino acids in most genomes (Fig. 4 and Table S6). However, tRNAs genes decoding histidine (tRNA^His^) were overall low abundant and even absent in 15 genomes (*A. domesticum*, *A. montevideense*, *A. porosum*, *Cutaneotrichosporon arboriformis*, *Cu. curvatus*, *Cu. cyanovorans*, *Cu. dermatis*, *P. fragicola*, *Trichosporon asahii* and *Tr. inkin*), while the aspartate encoding tRNA (tRNA^Asp^) was absent in *Pascua* and *Takashimella* spp. Additionally, some low copy number genes were exclusive to some species (e.g. Ala-GGC present only in *Ap. gamsii*) or clades (e.g. Ser-ACU was only detected in one sub-clade of *Cutaneotrichosporon* species). Interestingly, we observed that only one tRNA gene family per amino acid was expanded across different species (Fig. 4 and Table S6).

**Figure 4.**
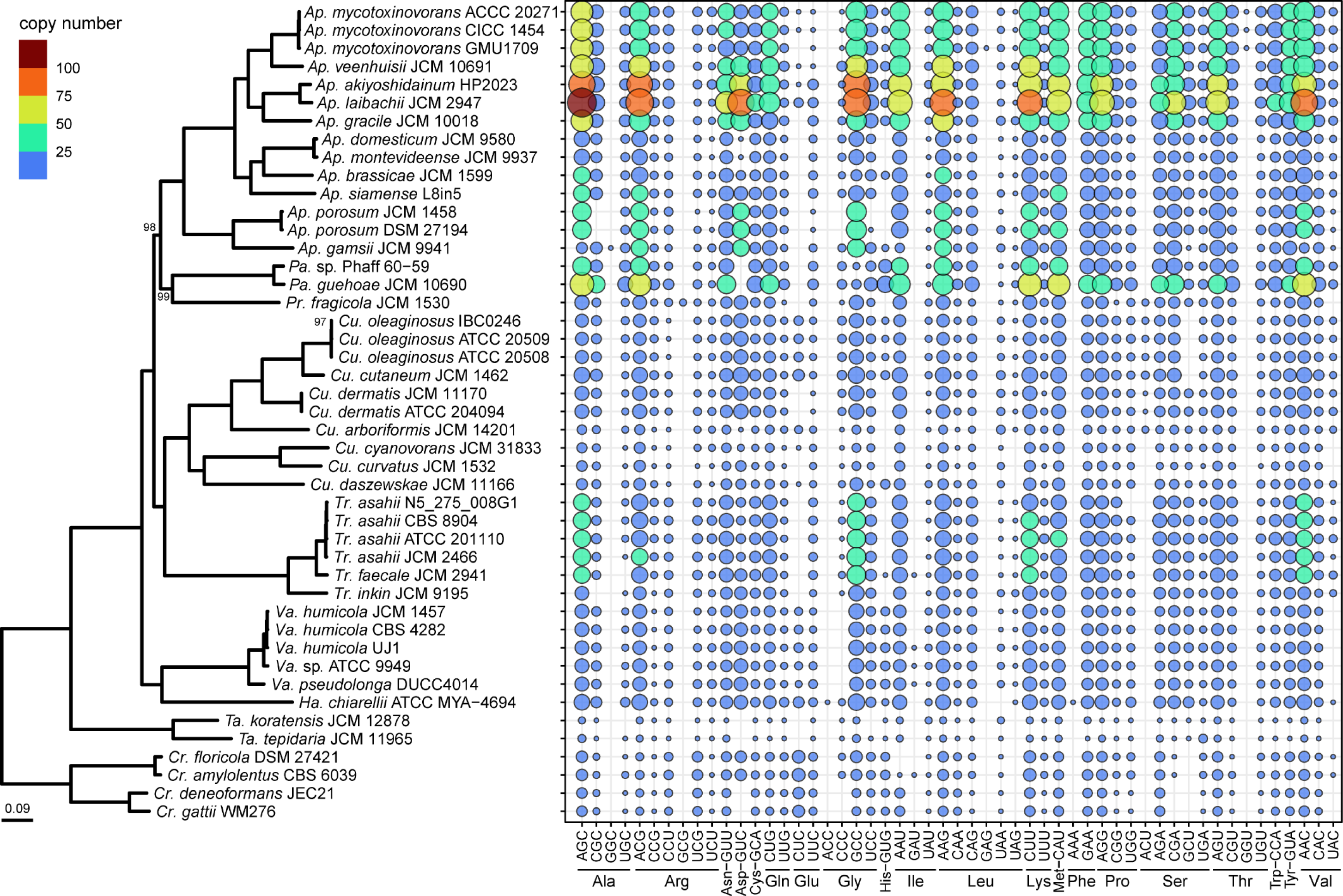
Distribution of tRNA gene copy number by anticodon and respective amino acid. Circle shapes are scaled and colour coded for gene copy number. Genera names are abbreviated (Ap., Apiotrichum; Pa., Pascua; Pr. Prillingera; Cu., Cutaneotrichosporon; Tr., Trichosporon; Va., Vanrija; Ha., Haglerozyma; Ta., Takashimella; Cr., Cryptococcus).

To understand if the evolutionary process of the tRNA expansion correlated with the phylogenetic relationship of species, we used the Blomberg’s K measure to assess evolution of the trait (i.e. in this case gene copy number) considering phylogenetic distance as the only predictor of trait similarity among species. Additionally, we tested the pairwise similarity and correlation of gene composition among species, considering their phylogenetic distance. For this, we calculated the pairwise Bray-Curtis similarity among all genomes, considering the copy number of each tRNA gene family gene. The pairwise branch length among all genomes was used as a measure of phylogenetic distance.

The tRNA gene composition (i.e. copy number per gene family) of 45 out of 55 tRNA genes produced a statistically significant phylogenetic signal (Blomberg’s *P*-value < 0.05). Most genes (38 out of 45) exhibited a low phylogenetic signal (Blomberg’s K < 1, Blomberg’s *P*-value < 0.05) (Fig. 5), indicating convergent evolution, since closely related species are less similar than expected in comparison to distantly related species, under a Brownian motion model of trait evolution. Only 5 genes showed a high phylogenetic signal (Blomberg’s K > 1, Blomberg’s *P*-value < 0.05), indicating stabilizing selection or low rates of evolution related to gene copy number, suggesting that this trait is highly conserved phylogenetically and closely related species are more similar than expected based on phylogenetic relationships. The lack of a significant phylogenetic signal (Blomberg’s *P*-value > 0.05) indicates that the remaining two genes evolved independently of the phylogenetic structure or due to reduced occurrence in the phylogeny. Furthermore, tRNA gene composition was strongly correlated and highly similar (Fig. 6) among distantly related species. Altogether, these analyses revealed the tRNA gene expansion, copy number and composition to be independent from the phylogenetic structure. This also indicates a dynamic tRNA repertoire where tRNA gene expansion occurred multiple and independent times in this group of species.

**Figure 5.**
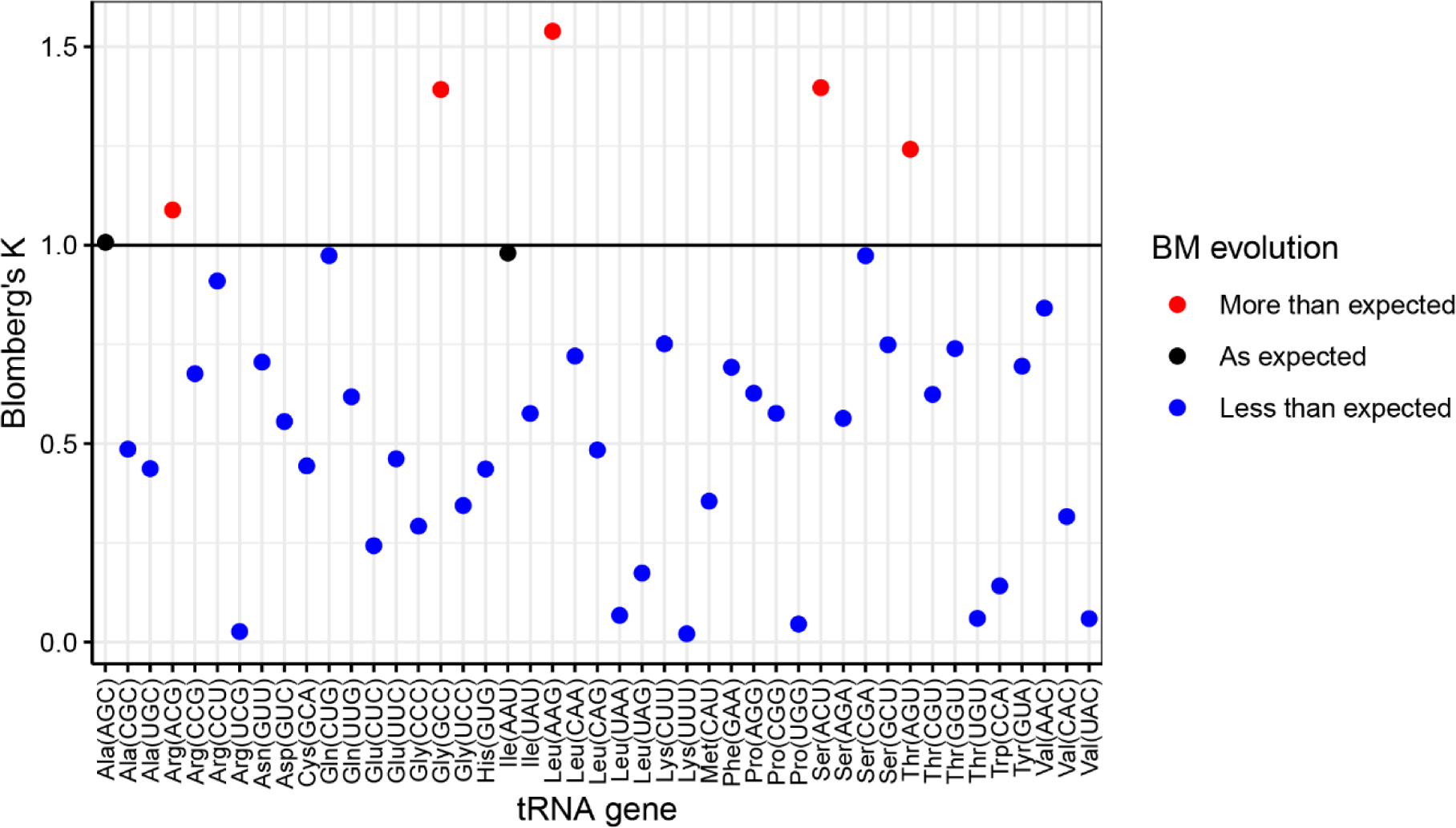
Variation in the phylogenetic signal (Blomberg’s K) of tRNA gene copy number in Trichosporonales. A total of 43/45 tRNA genes comprised Blomberg’s K not equal to 1 (± 0.02), suggesting they expanded less or greater than expected by genetic drift according to the Brownian Motion model of evolution. All signals are statistically significant (*P* <0.05).

**Figure 6.**
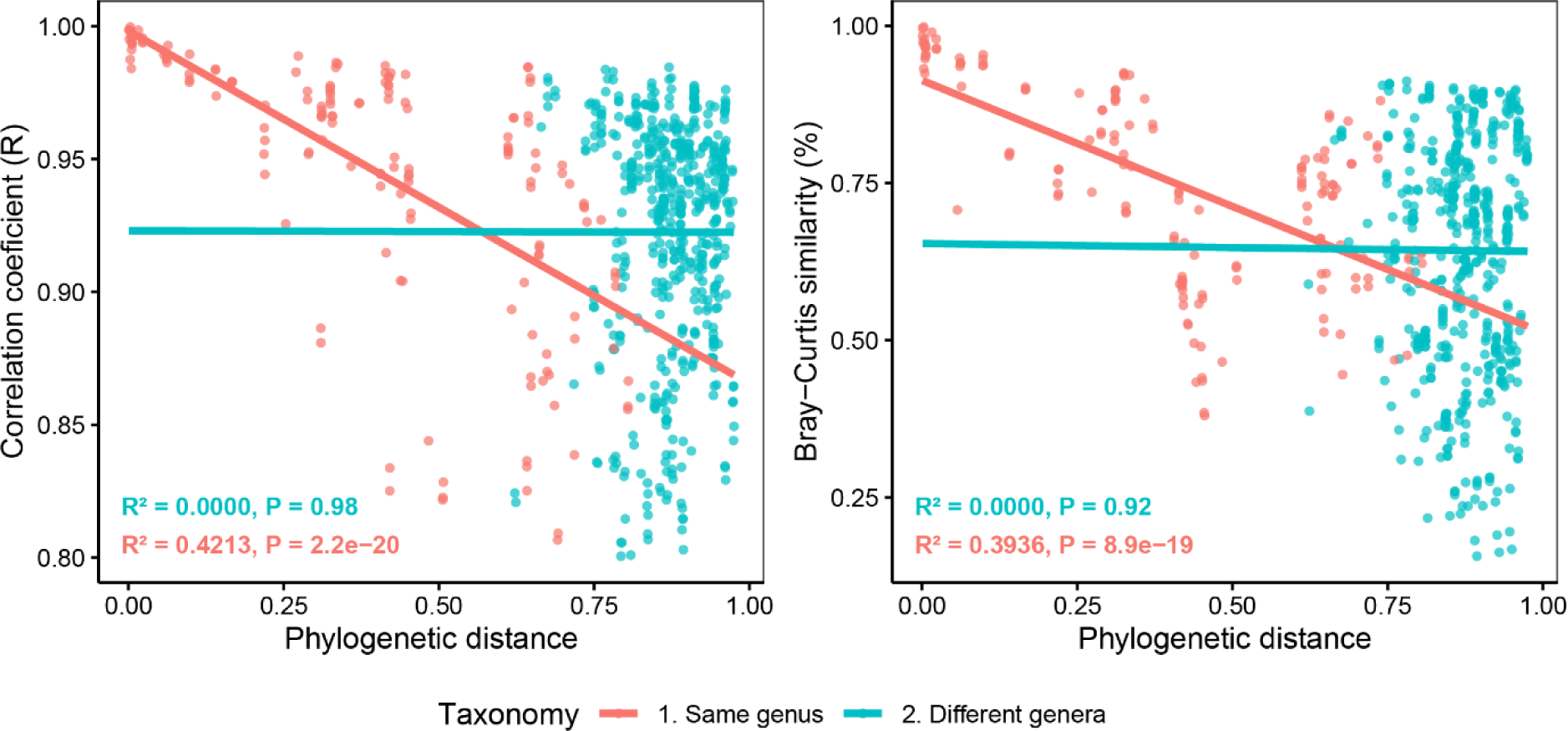
tRNA composition per taxonomic level in the Trichosporonales order. The pairwise correlation coefficient of individual gene copy number and Bray-Curtis similarity in relation to the respective phylogenetic distance is indicated for members of the same genus or different genera. Each data point represents one tRNA gene family. Due to their basal phylogenetic position, *Takashimella* species were excluded from this analysis.

### Intragenomic tRNA gene sequence variation suggests recent and multiple gene duplication events

To gain a better understanding of the observed tRNA expansion (Fig. 1, Fig. 4), we further assessed the intragenomic sequence variation of multiple-copy tRNA genes. Additionally, sequence variation among tRNA gene families was used to provide further insight into selection acting on these genes, as well as information about the evolutionary history of gene copies within each genome.

First, we investigated the relevance of gene copy-number on the variation among copies within genomes, for each gene family (i.e. genes with the same anticodon sequence). Here, we expected to find two possible scenarios: (i) genes with high copy-number to show a high level of conservation, suggesting recent gene duplication events, while genes with low copy-number to be more variable, suggesting ancient duplications (supporting Fig. 5 and Fig. 6); or (ii) high conservation of low copy genes due to high functional selective pressure, while relaxed selection on high-copy number genes due to functional gene redundancy.

Along Trichosporonales, the nucleotide variation (transitional and transversional substitution rates based on Kimura’s 2-parameter distance) in each tRNA gene family was inversely proportional to gene copy number (Fig. S7A). tRNA genes with a high copy number showed a low sequence variation, while genes with low copy number displayed the highest intragenomic variation (Fig. S7B). Variable positions among gene copies were mainly located in the predicted intronic region, but nucleotide substitutions and insertion/deletions were also detected in the tRNA body (data not shown). These results support the hypothesis of a recent gene expansion for the highly abundant tRNA gene families.

Next, we compared the nucleotide variation among all tRNA gene families within each genome to assess if some tRNAs exhibit more variation than others. Hereby we find that distinct tRNA genes decoding the same amino acid display different levels of variation. When amino acids were decoded by multiple genes, one of the genes displayed a higher level of conservation than the others (e.g. tRNA^Arg^-ACG and tRNA^Arg^-UCU) (Fig. S8). The tRNA with the highest intragenomic sequence variation was for the tRNA^Leu^-UAG gene (median distance: 0.76), while tRNA^Gly^-GCC was the most conserved gene (median distance: 0.02) (Fig. S8 and S9). These results suggest a differential selective pressure among genes decoding the same amino acid and also across genes decoding different amino acids.

### Interspecific tRNA gene sequence variation reveals different selective patterns

To address the selective pressure acting on tRNA genes across the phylogeny, we compared the overall tRNA nucleotide variation among all members of Trichosporonales.

Different species depicted different levels of nucleotide sequence variation in their tRNA genes (Fig. S10). The highest rate of tRNA sequence variation was detected in *Takashimella koratensis* (median distance of 0.32 per tRNA gene), while *Cutaneotrichosporon curvatus* exhibited the least variation (median distance of 0.06 per tRNA gene). No statistical significance was detected between gene within-species variation among tRNA copies and species lifestyles (*P* ≥ 0.05). These results suggest different selective pressure for each species, independently of the phylogenetic structure and lifestyle. Furthermore, this could also indicate that multiple and independent gene duplication events happened throughout the evolutionary history of this order, some being very recent.

### Translation efficiency of metabolic pathways is a predictor of lifestyle

Trichosporonales species are found in the environment and as opportunistic pathogens colonizing the skin or inner organs of the human hosts. To understand the effect of tRNA expansion and adaptation to lifestyle, we estimated the translation efficiency for genes known to be particularly relevant for certain lifestyles. Hereby, we focused on carbohydrate and lipid metabolic pathways, essential for growth in the environment (i.e. as a saprotroph), and in association with a human host (i.e. as an opportunistic pathogen), respectively.

Overall, we find that, in both Trichosporonales and Tremellales (Cryptococcus), all species consistently comprised an overall higher gene content related to carbohydrate transport and metabolism compared to genes related to lipid transport and metabolism (Fig. 7A). The copy number of genes related to lipid metabolism was generally higher in saprotrophic (median 197 genes) than for pathogenic (median 176.5 genes) *Cryptococcus* spp., however, this pattern was not consistent for *Apiotrichum* or *Cutaneotrichosporon* species (Fig. 7A). In the latter genera, opportunistic pathogenic species showed a higher content of genes related to lipid metabolism, with a few saprotrophic outliers showing a higher gene content (Fig. S11). These results do not support our initial hypothesis (i) that opportunistic pathogens have a higher diversity and copy number of genes associated with lipid transport and metabolism. An additional example, is the saprotrophic Pascua showing the highest lipid-related gene content among all genera (median 278 genes) (Fig. S11). Altogether, this suggests that an increased number of genes involved in lipid transport and metabolism is not a hallmark of opportunistic human pathogens.

**Figure 7.**
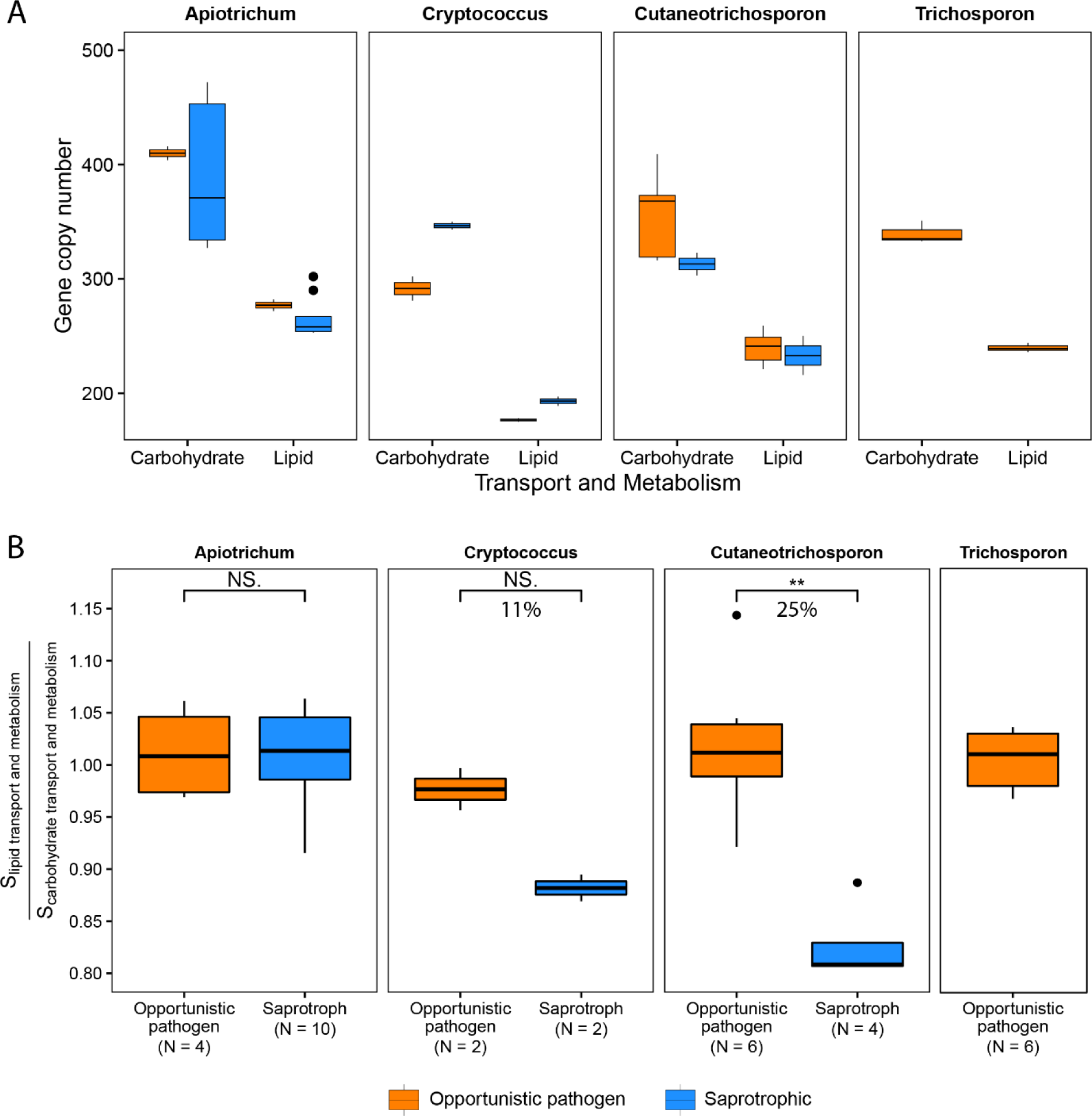
Gene copy number (A) and relative translation efficiency (B) for genes involved in lipid and carbohydrate transport and metabolism. Gene copy number and respective translation efficiency (S index) are compared within genera comprising both opportunistic pathogenic and saprotrophic species. Percentages indicate the increment of the S ratio between opportunistic pathogens and saprotrophic species.

To determine translational adaptation for genes related to lipid and carbohydrate metabolism, we predicted their codon optimization (S) for each pathway. Furthermore, we considered their individual response to lipids and carbohydrates as the ratio between the S of gene sets predicted to be involved in the transport and metabolism in these two metabolic pathways (S_lipid transport and metabolism_ : S_carbohydrate transport and metabolism_). This ratio was used to address different levels of optimization between genomes and compare the relative adaptive translation and efficiency of these metabolic pathways within each genome. This ratio indicates a similar level of adaptation to both lipid and carbohydrates (ratio ≈ 1), or a selective translational optimization towards of genes involved in carbohydrate (ratio < 1) or lipid (ratio > 1) transport and metabolism.

All opportunistic pathogens (*Apiotrichum* spp., *Cryptococcus* spp., *Cutaneotrichosporon* spp. and *Trichosporon* spp.) showed a similar optimization towards both lipid and carbohydrate substrates (S ratio ≈ 1) (Fig. 7B). Additionally, opportunistic pathogenic species showed an overall increased codon optimization (i.e. translation efficiency), in comparison to saprotrophic species, in genes involved in lipid transport and metabolism in relation to genes related to carbohydrate transport and metabolism (Fig. 7B). This pattern was consistent among *Cryptococcus* and *Cutaneotrichosporon* species. The codon optimization for lipid metabolism in relation to carbohydrate metabolism was 25% and 11% higher in opportunistic pathogenic *Cutaneotrichosporon* and *Cryptococcus* species, respectively, than in saprotrophic species of the respective genus. This indicates a higher translational adaptation of lipid-related metabolic genes in pathogenic species. Saprotrophic and pathogenic *Apiotrichum* species, however, showed a similar translation efficiency between lipid and carbohydrate pathways (S ratio ≈ 1). The measure of codon optimization was the highest in *Pascua* (S ratio > 1), while *Haglerozyma*, *Prillingera*, and *Vanrija* showed a similar optimization level to the pathogenic *Trichosporon*, *Cutaneotrichosporon* and *Apiotrichum* species (S ratio ≈ 1) (Fig. S12). Saprotrophic *Cryptococcus*, *Cutaneotrichosporon* and *Takashimella* showed consistently the lowest ratio of optimization, indicating a higher adaptation to carbohydrates than to lipid-rich substrates (S ratio < 1). These results support our hypotheses that lifestyle adaptations are imprinted in the codon optimization pathways relevant to lifestyles (iii) and that these are independent and convergent evolutionary events (v).

In summary, we determined that opportunistic pathogens are similarly adapted to both lipid- and carbohydrate-rich environments, based on their translational optimization of these metabolic pathways. Additionally, we found that saprotrophic *Cryptococcus* and *Cutaneotrichosporon* species without a pathogenic occurrence, have a higher codon optimization of carbohydrate-related gene, in comparison to lipids. However, saprotrophic Apiotrichum species are indistinguishable from other opportunistic pathogens, suggesting that these might represent potential pathogens. Our results suggest that translational optimization is an adaptive trait which may allow fungi to meet nutritional requirements under different conditions.

## Discussion

In this study, we inferred a robust phylogeny of the Trichosporonales and combined barcode data with phylogenomics to address species diversity and boundaries in this order. Our study provides the first comprehensive overview of the genome structure and tRNA diversity, evolution and respective implications on the emergence of pathogenicity in Tremellomycetes, including several emerging pathogens. Evidence suggests that pathogenicity emerged multiple times and independently. The emergence of pathogenicity is suggested to be linked to translational adaptation to animal hosts based on convergent tRNA gene expansion and codon optimization.

### Species and lifestyle diversity in Trichosporonales

The distribution of opportunistic pathogenic species across the phylogeny (Fig. 1) suggests that pathogenicity have emerged multiple times in the Tremellomycetes and specifically in the evolutionary history of the Trichosporonales and Tremellales orders. Detection of hybrid genomes belonging to common pathogenic species (Fig. S1) supports the hypothesis that emergence and speciation of fungal pathogens is often associated with hybridization (Stukenbrock 2013). The genomic variation in combination with the highly diverse ecology suggests that this order might possess an unreported high species diversity. Therefore, comprehensive taxonomic studies are required to clarify the fungal diversity in this group. Such studies are crucial for epidemiological studies and clinical settings, since closely related species commonly have different virulence and susceptibility to antifungal drugs (Hagen et al. 2015).

Species identification based on genomic data is often challenging due to the lack of reference genomes, mislabelled strains, or variable quality of databases (Houbraken et al. 2021). The use of genome-related indices, such as average nucleotide identity across the whole genome, is a powerful method to confirm species identification and determine species boundaries (Libkind et al. 2020; Houbraken et al. 2021). However, this approach is currently hindered by the shortage of genomic data. Therefore, the use of molecular barcodes, such as ITS and LSU regions, is an essential step for species identification and correct interpretation of the genomic data.

The rapid and accurate species identification is vital on medical settings to provide timely and appropriate treatment to patients. In our study, the pairwise comparison of genome averaged nucleotide identity (ANI) among all taxa suggests that the species boundaries are delimited by 97% similarity threshold (Table S3, Table S4 and Table S5). While species identification based on molecular barcodes is reliable in this group, genome ANI is a more effective method for species delimitation than based exclusively on ITS and LSU (Table S3, Table S4 and Table S5). However, there is a need to develop and implement more suitable barcodes for a more robust identification of fungal pathogens in clinical settings (Takashima et al. 2015). The use of a quicker and more accurate pathogen identification method will greatly enhance the efficiency and effectiveness of fungal infection diagnosis and treatment during medical emergencies.

### Convergent tRNA gene expansion and evolution

As reported for other fungi, other eukaryotes, and bacteria (Marck and Grosjean 2002; Chen et al. 2012; Wint et al. 2022) genomes of Trichosporonales species only comprise a subset of the full theoretical set of 61 tRNA gene families (Figs. 4 and S6, Table S6). The total number of distinct tRNA gene families (i.e. anticodon types) reported in previous studies correspond to the numbers that we predict for Trichosporonales (between 41 and 46). This is above the theoretical minimum of 30 anticodons for decoding the standard genetic code (Marck and Grosjean 2002). Interestingly, in our study, some codons and amino acids detected in protein-coding genes were missing complementary tRNAs in the same genome (Fig. 4, Table S6). Previous studies report that codon usage and codon abundance in protein-coding genes do not always correspond to the most abundant associated tRNA gene and some tRNAs may even remain undetected in the genome (Wilken et al. 2020; Chen et al. 2012). It has been suggested that, in fungi, tRNA wobbling of the third base position might enable a tRNA to recognize several unspecific synonymous codons and thereby allow the decoding of these amino acids (Chen et al. 2012; Wilken et al. 2020; Cannarozzi et al. 2010).

Studies report that some non-coding RNAs have regulatory roles which are independent of their primary functions (e.g. tRNA, mRNA) (Huang et al. 2021; Rudinger-Thirion et al. 2011; Geslain and Pan 2011). Variable tRNA gene sequences are also hypothesised to confer unique roles, but currently these remain to be determined (Goodenbour and Pan 2006). Similar to reports in other eukaryotes and fungi (Goodenbour and Pan 2006), we identified a high intragenomic sequence variation for tRNA genes having the same anticodon sequence (Figs. S7, S8 and S9). This variation was predominant in low-copy tRNA gene families. Evolutionary experiments in *Saccharomyces cerevisiae*, revealed that the tRNA pool within the genome can rapidly evolve to meet translational demands (Yona et al. 2013). Furthermore, mutations in the tRNA genes are also a common adaptive mechanism to obtain a higher fitness when presented with new environmental challenges (Yona et al. 2013). Our findings highlight that the evolution of individual tRNA genes families is characterized by different selective pressures. The increase in the tRNA gene copy number is thought to be the result of genome expansion (Santos and Del-Bem 2023), which is also supported here by our results (Fig. S5). In some species tRNA expansion has been linked directly with lifestyle transitions such as in the Sodariomycetes, where tRNA expansion has been associated with pathogenic lifestyles (Fijarczyk et al. 2022). tRNA expansions are also shown to be a response to changes in codon usage (McDonald et al. 2015), which, as found here, could imply adaptation of species to new ecological niches, namely host-association. tRNA gene content is a key genomic feature that contributes to the translational efficiency and adaptation, but the mechanisms driving the tRNA gene copy number and codon composition are currently poorly understood (Novoa et al. 2012). Analyses combining phylogenetic information and tRNA numbers, indicate that the expansion of these genes is independent and possibly a result of convergent evolution (Fig. 5 and Fig. 6). Furthermore, tRNA gene composition among species is not fully explained by phylogenetic relatedness that is expected under genetic drift (Kamilar and Cooper 2013; Wint et al. 2022). Correlative and similarity analyses based on the structure of the tRNA pool (i.e. relative abundance of each gene family) supports independent and convergent evolutionary expansion of tRNA genes along the phylogeny. Convergent evolution is generally attributed to shared molecular, genetic, physiological, or ecological constraints, which can limit or bias the genomic or phenotypic variation among organisms (Losos 2011). Alternatively, convergent evolution can also be a result of stochastic processes, albeit to a lesser extent (Losos 2011). The convergence of codon usage frequencies and optimization at a macroevolutionary level may be an indication of shared constraints, which are imposed by both neutral and adaptive pressures (Wint et al. 2022). Further research is needed to clarify the mechanisms driving tRNA genes copy number and codon composition during evolutionary processes. This knowledge may provide insights into the adaptive mechanisms that influence tRNA expansion and convergence, while contributing to our understanding on how organisms adapt to new ecological niches.

### Adaptive translation and pathogenicity in Trichosporonales

Genome evolution and translational regulation can be linked to pathogenicity and the ability of pathogenic species to colonize multiple hosts (Badet et al. 2017). Comparative genomic approaches suggest that several saprotrophic species share many features with those reported as opportunistic pathogens (Fig. 3). Several saprotrophic species (Apiotrichum, Haglerozyma, Pascua, Prillingera and Vanrija) show a similar adaptation to lipids as others reported as dermatophytes and opportunistic pathogens (Apiotrichum, Cutaneotrichosporon, Trichosporon and Cryptococcus) (Figs. 7, S11 and S12). This raises the concern that some species currently classified as saprotrophic are potentially opportunistic pathogens. During evolution, tRNA genes and codon composition coevolve to tune the expression of individual genes in relation to the whole proteome (Frumkin et al. 2018). Additionally, genes required under different environmental conditions, contain a codon bias similar to highly expressed genes (Parvathy et al. 2022). Opportunistic pathogenic fungal species are currently classified based on human clinical reports.

Several Trichosporonales species have been isolated as commensals from the skin of healthy giant Pandas (*Ailuropoda melanoleuca*) (Ma et al. 2019). These species included *Apiotrichum laibachii*, *Ap. brassicae*, *Ap. gracile*, *Ap. domesticum*, *Pascua guehoae*, *Trichosporon asteroides*, *Cutaneotrichosporon jirovecii*, *Cu. cutaneum*, *Cu. moniliiforme*, *Cu. spelunceum*, and *Cu. middelhovenii*. Pathogenicity tests in mice revealed that all of these isolates caused successful skin and invasive infections in immunosuppressed mice (Ma et al. 2019). Except for *Ap. domesticum*, *Ap. gracile* and *Cu. moniliiforme*, all species caused significant damage to the liver and skin of healthy immunocompetent mice (Ma et al. 2019). The authors also report that different species and genera caused similar pathological lesions on the skin and liver. Additional animal infections have been reported for *Ap. domesticum* in a urinary infection in a cat (Sakamoto et al. 2001) and for *Ap. montevideense* in a meningoencephalitis in a dog (Bryan et al. 2014). The human opportunistic pathogen *Tr. asahii* is capable of infecting immunocompetent mice and disseminate into their kidneys, liver, spleen and brain (Montoya et al. 2018). In addition to *Tr. asahii*, *Tr. asteroides* and *Tr. inkin* are able to successfully infect immunocompromised mice and larvae of *Galleria mellonella*, inducing a high mortality rate on the infected subjects (Mariné et al. 2015). These studies and reports provide evidence of the pathogenic potential of species currently classified as saprotrophic. This further supports our hypotheses that the saprotrophic *Apiotrichum* and *Pascua* species could represent potential human opportunistic pathogens, as suggested by their adapted translation to the metabolism of lipids which is comparable to that of other pathogens (Figs. 7, S11 and S12). Infection mechanisms appear to be shared among the different genera, as pathological characteristics on mice are similar when infected by different species and genera (Ma et al. 2019). Pathogenicity tests and clinical reports also indicates that this order could have a broad range of hosts, including mammals (humans, mice, dogs, cats) and insects.

The colonization of new host species (i.e. host jump or host shift), switches in lifestyle and their importance for pathogenicity, are best studied events on plant pathogenic fungi. Emergence and speciation of fungal plant pathogens have been suggested to be associated with host domestication, host jumps and hybridization (Stukenbrock 2013). Interestingly, in Trichosporonales, interspecies hybrid genomes (i.e. hybrid individuals) have been predicted for the human pathogenic species *Cutaneotrichosporon dermatis* (Fig. S1) and reported for *Trichosporon coremiiforme*, *Trichosporon ovoides* and *Cutaneotrichosporon mucoides* (Takashima et al. 2018). This suggests recent and/or ongoing adaptation events in this group of fungi, which might be related with the emergence of pathogenicity among Trichosporonales. Accordingly, future comparative genomic studies, might focus on predicting host ranges and evolutionary events leading to the emergence of pathogenicity in Trichosporonales.

The similarities observed in many genomic features between saprotrophic species and opportunistic pathogens raise concerns about the potential of some saprotrophic species to easily become opportunistic human pathogens. This urges the development of tools to accurately classify species based on their ecological strategies by using genomic and physiological data, as using only species occurrence data to classify lifestyles may not be sufficient. Comparative genomic studies can be used to predict future fungal infection outbreaks by identifying species with similar genomic signatures to known pathogens. Furthermore, the coevolution of tRNA genes and codon composition can be linked to the translational regulation of genes under different environmental conditions (Parvathy et al. 2022), which may have implications for the ability of pathogenic species to colonize different hosts (Badet et al. 2017). A deeper understanding of evolutionary processes associated with lifestyle adaptation is necessary to develop effective strategies to control fungal infections.

### Conclusions

In this study, our analyses link genomic information with ecology and fungal lifestyle across the Tremellomycetes. Our results indicate that the pathogenic lifestyle evolved several times throughout the evolutionary history of the Tremellales and Trichosporonales. We detected independent and convergent expansion of the translation machinery, while detecting lifestyle adaptation signatures in the translation efficiency of metabolic genes related to lifestyle. We find evidence for an evolutionary scenario where distinct habitats select for an optimized translation of genes involved in successful colonization of the respective habitat. Our results also suggest that since saprotrophic species are indistinguishable from opportunistic pathogens based on many genomic features, these fungi might have the potential to easily evolve towards pathogenic lifestyles. We predict that lifestyles are not strictly defined by gene repertoires, but also by expression profiles and translation efficiency of relevant pathways. Our study provides basis for future research addressing evolutionary gene regulation, translational adaptation and dynamic lifestyles in fungi. In conclusion, our study highlights the complexity of evolutionary mechanisms defining fungal lifestyle. The functional implications of the reported adaptation patterns remain to be determined.

## Acknowledgements

We thank all the members of the Environmental Genomics (Stukenbrock’s lab) for support and helpful discussions. We thank the Max Planck Institute for Evolutionary Biology and the respective IT team for the computing infrastructure and technical support. This project has been funded by the Deutsche Forschungsgemeinschaft (DFG) Walter Benjamin Programme (Grant No. GU 2252/1-1, Project No. 460261834) awarded to MG.

## Declaration of competing interest

The authors declare that they have no competing interests.

## Notes

### Competing Interest Statement

The authors have declared no competing interest.

