## Supplementary information for "Lifestyle transitions in basidiomycetous fungi are reflected by tRNA composition and translation efficiency of metabolic genes"

**Table S1.** Isolates and genome assemblies analysed in this study and respective relevant metadata is provided. The status of the strain is indicated for type-strains (T) and hybrids (H). Accession numbers from NCBI and links for JGI and ATCC for the genome assemblies are provided. When available, accession numbers for additional genome assemblies for the same strain are provided between parentheses. Strain IDs used throughout the text and figures are highlighted in bold, while individuals further selected as species representatives for statistical analyses are underlined.

| Species | Strain/Voucher | Strain status | Assembly accession | Strain source | Genome source | Species clinical reports |
|---|---|---|---|---|---|---|
| **Trichosporonales** | | | | | | |
| *Apiotrichum akiyoshidainum* | **HP2023**, CBS 10550, ATCC MYA-4129 | | GCA_002973495.1 | [1] | [2] | NA |
| *Apiotrichum brassicae* | **JCM 1599**$^{T}$, CBS 6382$^{T}$, ATCC 24124$^{T}$, AJ 4833, BCRC 21783, IFO 1584, NBRC 1584, NCYC 2562, UAMH 7657, CCRC 21783 | T | GCA_001600295.1 | [3] | [4] | NA |
| *Apiotrichum domesticum* | **JCM 9580**$^{T}$, CBS 8280$^{T}$, NBRC 103893, NCYC 2665 | T | GCA_001599015.1 | [5] | [6] | NA |
| *Apiotrichum gamsii* | **JCM 9941**$^{T}$, CBS 8245$^{T}$, JCM 12197, BCRC 22274, IFM 49226, NBRC 103894 | T | GCA_001600315.1 | [7] | [4] | NA |
| *Apiotrichum gracile* | **JCM 10018**$^{T}$, CBS 8189$^{T}$, CBS 102.10, NBRC 103905, UAMH 7662 | T | GCA_001600335.1 | [8] | [4] | NA |
| *Apiotrichum laibachii* | **JCM 2947**$^{T}$, CBS 5790$^{T}$, CGMCC 2.1972, NBRC 102674, PYCC 3865, UAMH 7666 | T | GCA_001600735.1 | [8] | [4] | NA |
| *Apiotrichum montevideense* | **JCM 9937**$^{T}$, CBS 6721$^{T}$, IMUR 2329, NBRC 103899, NCYC 2638, NRRL Y-7944, UAMH 7669 | T | GCA_001598995.1 | [8, 9] | RIKEN BRC JCM | NA |
| *Apiotrichum mycotoxinivorans* | **CICC 1454**, AS 2.1374 | | GCA_013177335.1 | NA | [10] | [11, 12] |
| *Apiotrichum mycotoxinivorans* | **GMU1709** | | GCA_011290525.1 | [12] | [13] | |
| *Apiotrichum mycotoxinivorans* | **ACCC 20271**, CGMCC 2.1374 | | GCA_001613755.1 | NA | [14] | |
| *Apiotrichum porosum* | **JCM 1458**$^{T}$, CBS 2040$^{T}$, NRRL Y-17198, VKM Y-3 | T | GCA_001600255.1 | [15] | RIKEN BRC JCM | NA |
| *Apiotrichum porosum* | **DSM 27194**, TPST6 | | GCF_003942205.1 | [16] | [17] | |
| *Apiotrichum siamense* | **L8in5**, CLIB 3781 | | GCA_023653615.1 | [18] | [19] | NA |
| *Apiotrichum veenhuisii* | **JCM 10691**$^{T}$, CBS 7136$^{T}$ | T | GCA_001600595.1 | [20] | RIKEN BRC JCM | [21] |
| *Cutaneotrichosporon arboriformis* | **JCM 14201**$^{T}$, CBS 10441$^{T}$, IFM 54862 | T | GCA_002335565.1 | [22] | [23] | [22] |
| *Cutaneotrichosporon curvatus* | **JCM 1532**$^{T}$, CBS 570$^{T}$, ATCC 10567$^{T}$, DSM 101032$^{T}$, AJ 4763, DBVPG 6206, IFO 1159, NBRC 1159, NCYC 476, NRRL Y-1511, PYCC 2565, VKM Y-2230, MUCL 29819, SBUG-Y 855 | T | GCA_001600275.1 (GCA_020080345.1, GCA_001028165.1) | NA | [4] | [24, 25] |
| *Cutaneotrichosporon cutaneum* | **JCM 1462**$^{T}$, CBS 2466$^{T}$, ATCC 28592$^{T}$, DSM 27285$^{T}$, BCRC 21675, DBVPG 6973, IFO 1198, IGC 3462, NBRC 1198, NCYC 444, NRRL Y-1490, PYCC 3462, UAMH 7659, MUCL 30308 | T | GCA_001600715.1 | NA | [4] | [26] |

| Species | Strain/Voucher | Strain status | Assembly accession | Strain source | Genome source | Species clinical reports |
|---|---|---|---|---|---|---|
| *Cutaneotrichosporon cyanovorans* | **JCM 31833**$^{T}$, CBS 11948$^{T}$, NRRL Y-48730, UOFS Y-2244 | T | GCA_002335625.1 | [27] | [23] | NA |
| *Cutaneotrichosporon daszewskae* | **JCM 11166**$^{T}$, CBS 5123$^{T}$, MUCL 30649, NBRC 100670 | T | GCA_002335585.1 | [28] | [23] | [28] |
| *Cutaneotrichosporon dermatis* | **JCM 11170**$^{T}$, CBS 2043$^{T}$, NBRC 102675 | T | GCA_003116895.1 | [29] | [23] | [30, 31] |
| *Cutaneotrichosporon dermatis* | **ATCC 204094**, Vitek 303483, API 7704022 | | genomes.atcc.org/genomes/ e80264e2adb34f72 | NA | ATCC | |
| *Cutaneotrichosporon dermatis* | **NICC30027** | H | GCA_020011015.1 | [32] | [33] | |
| *Cutaneotrichosporon mucoides* | **JCM 9939**$^{T}$, CBS 7625$^{T}$, IP 151, UAMH 7670 | T, H | GCA_003116955.1 | NA | [23] | [31, 34] |
| *Cutaneotrichosporon oleaginosus* | **ATCC 20509**$^{T}$, DSM 11815$^{T}$ | T | genomes.atcc.org/genomes/ 1bd8cbf8d02b479c (GCA_001712445.1) | [35] | ATCC | NA |
| *Cutaneotrichosporon oleaginosus* | **ATCC 20508** | | GCA_008065305.1 | [35] | [36] | |
| *Cutaneotrichosporon oleaginosus* | **IBC0246** | | GCF_001027345.1 | NA | [37] | |
| *Haglerozyma chiarellii* | **ATCC MYA-4694**$^{T}$, JCM 31834$^{T}$, CBS 11177$^{T}$ | T | mycocosm.jgi.doe.gov/Trich1 | [38] | JGI | NA |
| *Pascua guehoae* | **JCM 10690**$^{T}$, CBS 8521$^{T}$ | T | GCA_001600415.1 | [39] | RIKEN BRC JCM | NA |
| *Pascua 'guehoae'* sp. | **Phaff 60-59** | | mycocosm.jgi.doe.gov/Trigue1 | NA | JGI | NA |
| *Prillingera fragicola* | **JCM 1530**$^{T}$, CBS 8898$^{T}$, AJ 4683, NBRC 100667 | T | GCA_002335605.1 | [28] | [23] | NA |
| *Trichosporon asahii* | **JCM 2466**$^{T}$, CBS 2479$^{T}$, IGC 3469, NBRC 103889, NCYC 2677, UAMH 7654 | T | GCA_001972365.1 (GCF_000293215.1) | NA | [4] | [31, 40] |
| *Trichosporon asahii* | **CBS 8904** | | GCA_000299215.2 | NA | [41] | |
| *Trichosporon asahii* | **N5_275_008G1** | | GCA_004026345.1 | [42] | [42] | |
| *Trichosporon asahii* | **ATCC 201110**, TIMM 1318 | | genomes.atcc.org/genomes/ b7621150fd7849ee | [43] | ATCC | |
| *Trichosporon coremiiforme* | **JCM 2938**$^{T}$, CBS 2482$^{T}$, PYCC 3472, UAMH 7658 | T, H | GCA_001752605.1 | NA | [4, 44] | [40] |
| *Trichosporon faecale* | **JCM 2941**$^{T}$, CBS 4828$^{T}$, IGC 3475, IMUR 932, NBRC 103892, PYCC 3475, UAMH 7661 | T | GCA_001752585.1 | NA | [44] | [40] |
| *Trichosporon inkin* | **JCM 9195**$^{T}$, CBS 5585$^{T}$, ATCC 18020$^{T}$, BCRC 21503, IFO 10131, IGC 3727, NBRC 10131, NRRL Y-7793 | T | GCA_001752625.1 (GCA_004023515.1) | NA | [44] | [31, 40] |
| *Trichosporon ovoides* | **JCM 9940**$^{T}$, CBS 7556$^{T}$, RV 34067, UAMH 7671 | T, H | GCA_001752645.1 | NA | [44] | [31] |
| *Trichosporon ovoides* | **2NF903A** | H | GCA_009833065.1 | NA | Heilongjiang bayi agricultural university | |

| Species | Strain/Voucher | Strain status | Assembly accession | Strain source | Genome source | Species clinical reports |
|---|---|---|---|---|---|---|
| *Vanrija humicola* | **JCM 1457**$^{T}$, CBS 571$^{T}$, ATCC 14438$^{T}$, BCRC 21639, DBVPG 6019, IFO 0760, IGC 3387, MUCL 29840, NBRC 0760, NCYC 818, NRRL Y-12944, PYCC 3387, VKM Y-2238 | T | GCA_001600235.1 | NA | [4] | NA |
| *Vanrija humicola* | **CBS 4282** | | GCA_008065275.1 | NA | [36] | |
| *Vanrija humicola* | **UJ1**, JCM 9575, MUCL 30690, NBRC 100671 | | GCA_002897395.1 | NA | [45] | |
| *Vanrija 'humicola'* sp. | **ATCC 9949**, JCM 1459, CBS 2041, DSM 6382, IAM 12184, IFO 0753, NBRC 0753, NRRL Y-1266 | | genomes.atcc.org/genomes/ 781d93df71954299 | NA | ATCC | NA |
| *Vanrija pseudolonga* | **DUCC4014** | | GCA_020906515.1 | [46] | Dankook University | NA |
| *Takashimella koratensis* | **JCM 12878**$^{T}$, CBS 10484$^{T}$, TY-242, TISTR 5816 | T | GCA_003116875.1 | [47] | [23] | NA |
| *Takashimella tepidaria* | **JCM 11965**$^{T}$, CBS 9427$^{T}$ | T | GCA_003116915.1 | [48] | [23] | NA |
| **Tremellales** | | | | | | |
| *Cryptococcus amylolentus* | **CBS 6039**$^{T}$, JCM 1690$^{T}$, ATCC 56469$^{T}$, IFO 10423, IGC 4486, NRRL Y-7784, PYCC 4486 | T | GCF_001720205.1 | NA | [49] | NA |
| *Cryptococcus deneoformans* | **JEC21**, CBS 10513, ATCC MYA-565, B-4500 | | GCF_000091045.1 | [50] | [51] | [52] |
| *Cryptococcus floricola* | **DSM 27421**$^{T}$ | T | GCA_006352305.1 | [53] | [53] | NA |
| *Cryptococcus gattii* | **WM276** | | GCA_000185945.1 | NA | [54] | [52] |

NA, not available; RIKEN BRC JCM: https://www.jcm.riken.jp/cgi-bin/jcm/jcm_genomelist; ATCC: https://genomes.atcc.org/genomes/mycology; JGI: https://mycocosm.jgi.doe.gov/mycocosm/home

**Table S2.** Pairwise comparisons of the barcode regions (ITS and LSU) obtained from whole genome sequencing and from the respective type-strain based on Sanger sequencing. The accession numbers for the publicly available sequences from the type-strain barcodes are provided. Single accession numbers contain both barcode regions. The similarity (%) and mismatches (number of nucleotide substitutions and insertions/deletions) between both datasets are indicated. Whole genome sequencing information is provided on Table S1. NA indicates that it was not possible to retrieve the barcodes from the corresponding genomic data.

**Table S3.** Genomic distance matrix among Trichosporonales genomes. The lower triangle contains pairwise the averaged nucleotide identity (ANI, %) between genomes, while the upper triangle contains the corresponding pairwise phylogenetic distance. The phylogenetic distance was calculated based on the maximum likelihood phylogeny (Fig. 1). Pairwise comparisons within species are highlighted.

**Table S4.** Barcode (ITS and LSU) distance matrix among Trichosporonales genomes. The lower triangle contains pairwise nucleotide substitutions and insertions/deletions, while the upper triangle contains pairwise sequence similarity. The overall alignment contained 1143 sites. Pairwise comparisons within species are highlighted. NA indicates that it was not possible to retrieve the barcodes from the corresponding genomic data.

**Table S5.** Summary of the phylogenetic analyses of selected taxa based on molecular barcodes (ITS and LSU regions) and genome averaged nucleotide identity (ANI). The complete pairwise comparisons are provided in Table S3 and Table S4.

| Taxa | Strain comparison | ITS/LSU similarity (mismatches) | Genome ANI (%) |
|---|---|---|---|
| *Apiotrichum domesticum* **x** *Apiotrichum montevideense* | JCM 9580$^{T}$ **x** JCM 9937$^{T}$ | 99.7% (3 bp) | 92.11% |
| *Apiotrichum mycotoxinivorans* | CICC 1454* **x** GMU1709 **x** ACCC 20271 | 100.0% (0 bp) | 98.89% – 99.93% |
| *Apiotrichum porosum* | DSM 27194 **x** JCM 1458$^{T}$ | 100.0% (0 bp) | 97.16% |
| *Cutaneotrichosporon dermatis* | JCM 11170$^{T}$ **x** ATCC 204094 | 99.9% (1 bp) | 99.97% |
| *Cutaneotrichosporon oleaginosus* | ATCC 20508 **x** IBC0246 **x** ATCC 20509$^{T}$ | 99.6% (4 bp) – 100.0% (0 bp) | 99.96% – 99.98% |
| *Pascua* sp. **x** *Pascua guehoae* | Phaff 60-59 **x** JCM 10690$^{T}$ | 99.5% (5 bp) | 87.81% |
| *Trichosporon asahii* | ATCC 201110 **x** CBS 8904* **x** JCM 2466$^{T}$ **x** N5_275_008G1* | 99.9% (1 bp) | 98.15% – 99.94% |
| *Vanrija humicola* | CBS 4282 **x** UJ1 **x** JCM 1457$^{T}$ | 100.0% (0 bp) | 97.77% – 98.62% |
| *Vanrija* sp. **x** *Vanrija humicola* | ATCC 9949 **x** JCM 1457$^{T}$ | 99.2% (8 bp) | 90.79% |

* No barcodes available

**Table S6.** Distribution of tRNA genes among Trichosporonales fungi on the genome assembly data. The overall number of detected genes is provided, as well as, by anticodon and the corresponding decoded number of amino acids. The total number of genes per decoded amino acid and anticodon is also provided.

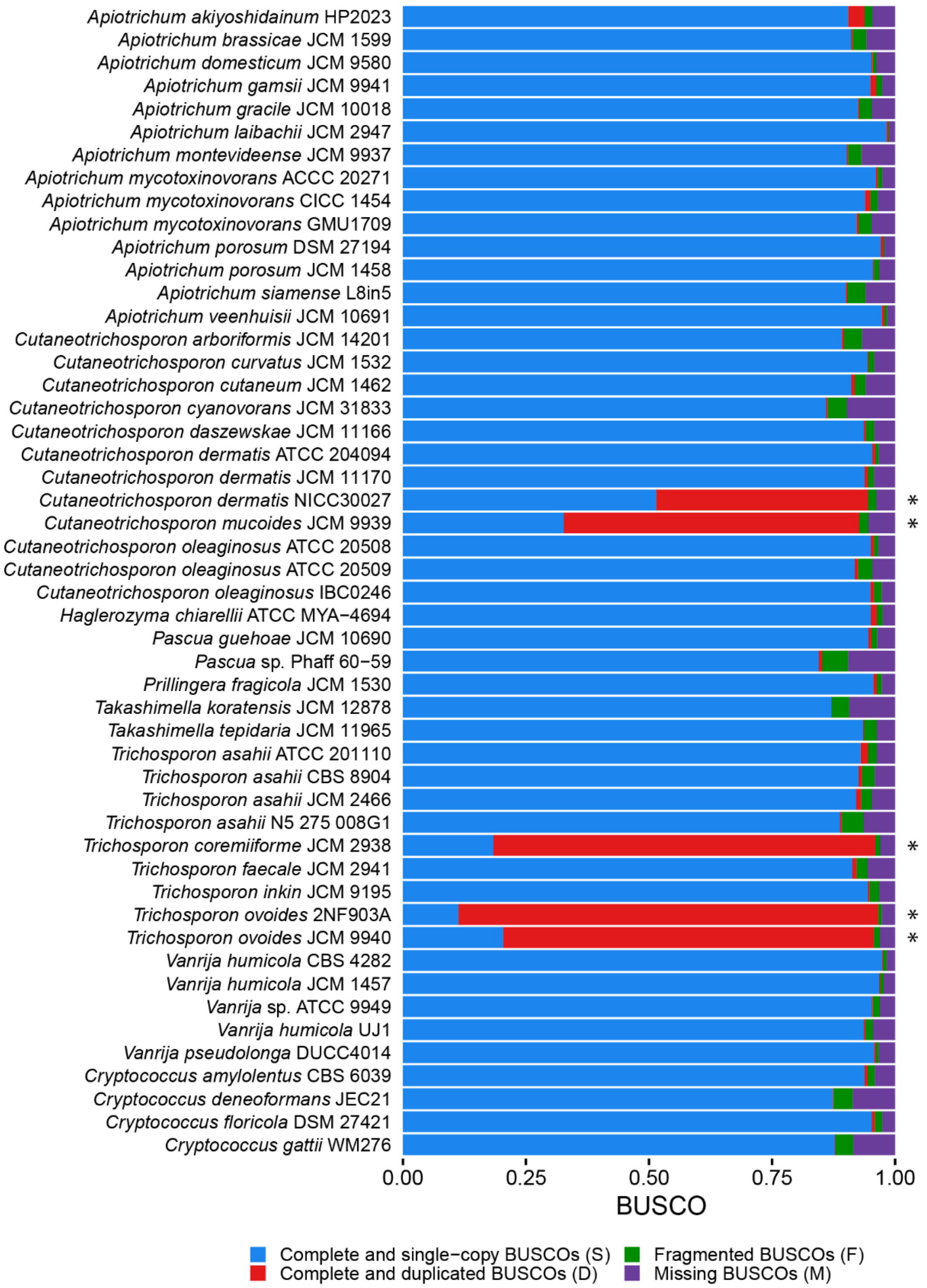


**Figure S1.** Assessment of proteome completeness. Asterisks indicate genome assemblies that, due to high content of duplicated BUSCOs, are suspected to be hybrids or diploid genomes and were therefore excluded from further analyses. BUSCO analyses were performed on predicted proteomes.

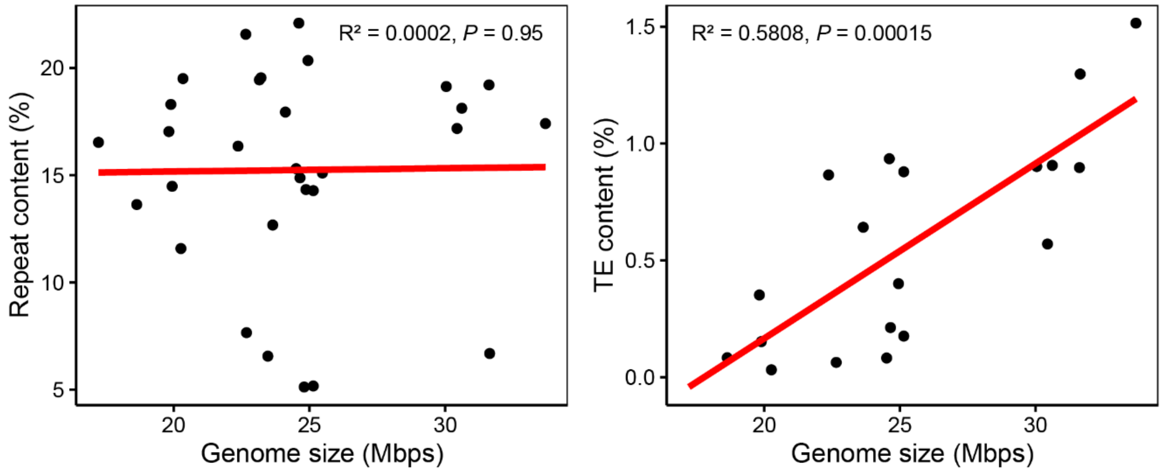


**Figure S2.** Correlations between genome size, repeat content and transposable element (TE) content. Linear regressions (red line) and respective correlations ($R^2$ and *P*-value) are indicated.

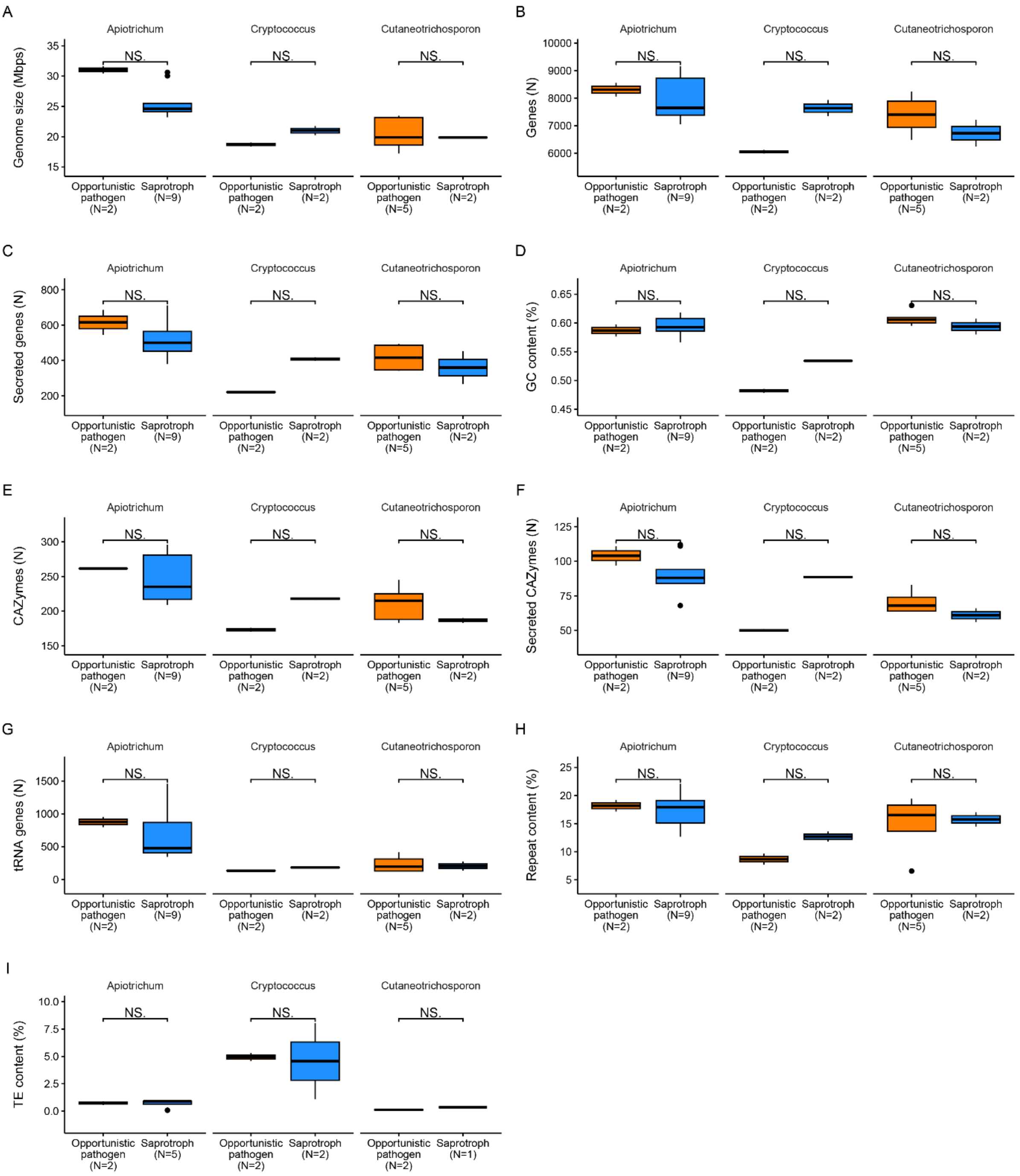


**Figure S3.** Comparison of genomic features between opportunistic pathogenic and saprotrophic lifestyles within Apiotrichum, Cryptococcus and Cutaneotrichosporon genera. No significant statistical differences ("NS." *P*-value ≥ 0.05) were observed based on Wilcoxon signed-rank test due to low number of samples. To avoid overestimation, only one genome per species was considered and the total number of considered strains is indicated.

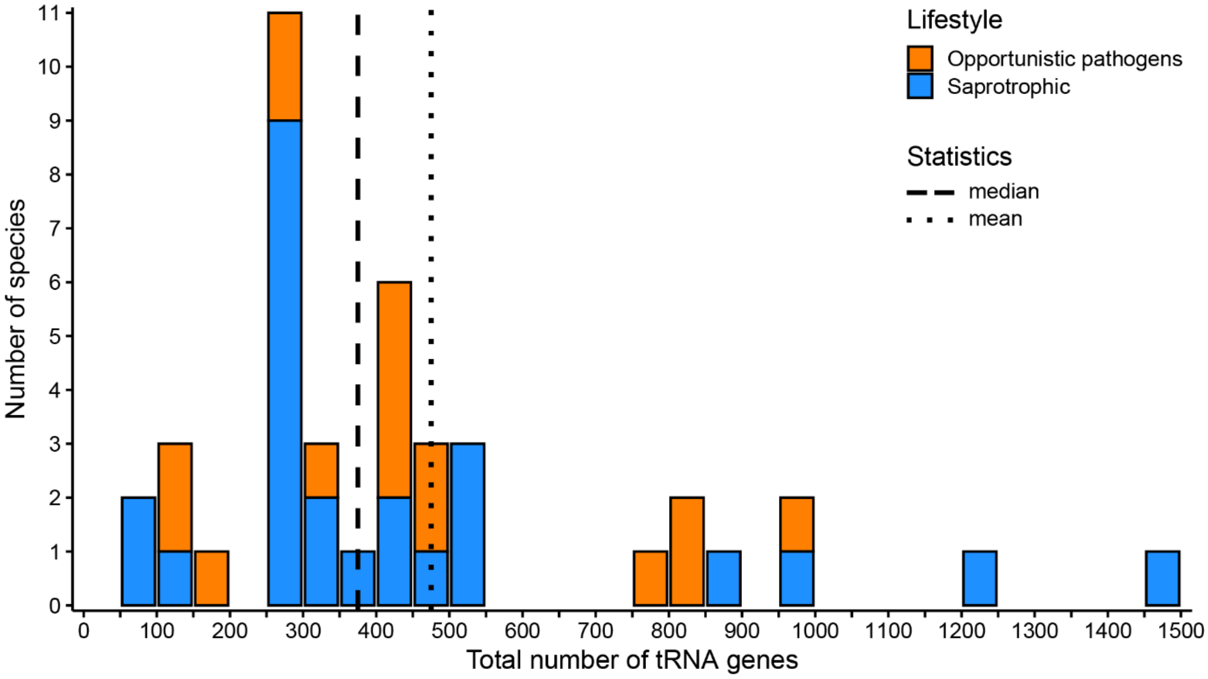


**Figure S4**. Distribution of the total number of tRNA genes across 41 fungal genomes the Trichosporonales order.

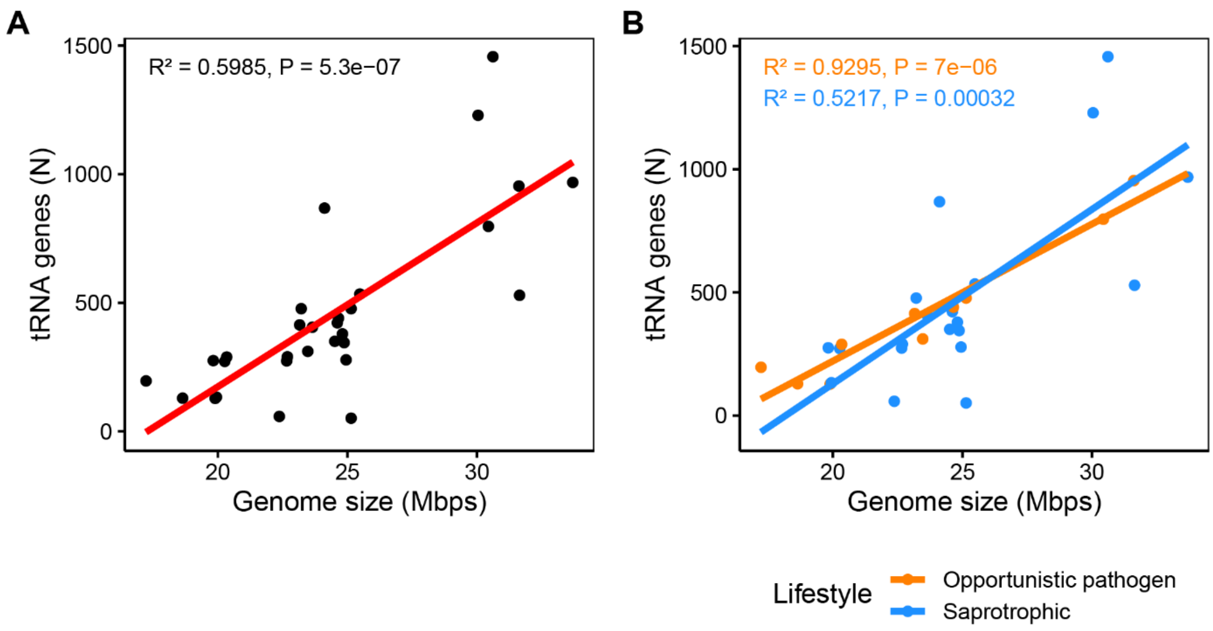


**Figure S5.** Correlations between genome size and number of tRNA genes. Linear regressions and respective correlations are indicated for all Trichosporonales (A) or by lifestyle (B).

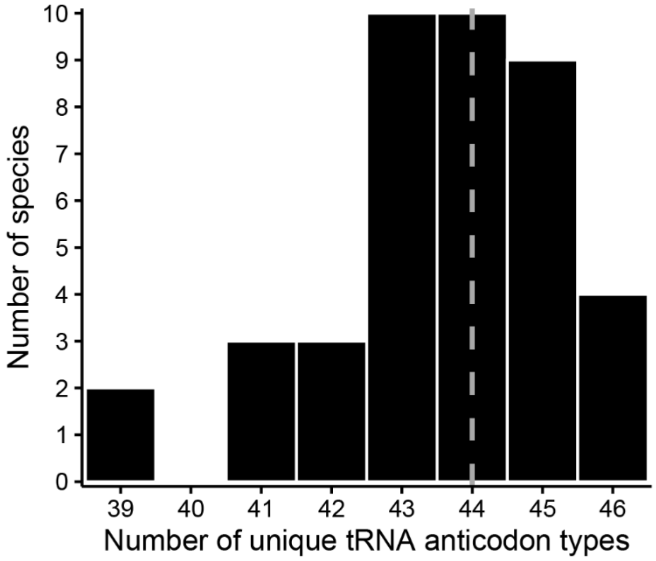


**Figure S6.** Distribution of unique tRNA anticodon types among Trichosporonales species. Dotted grey line represents the median value.

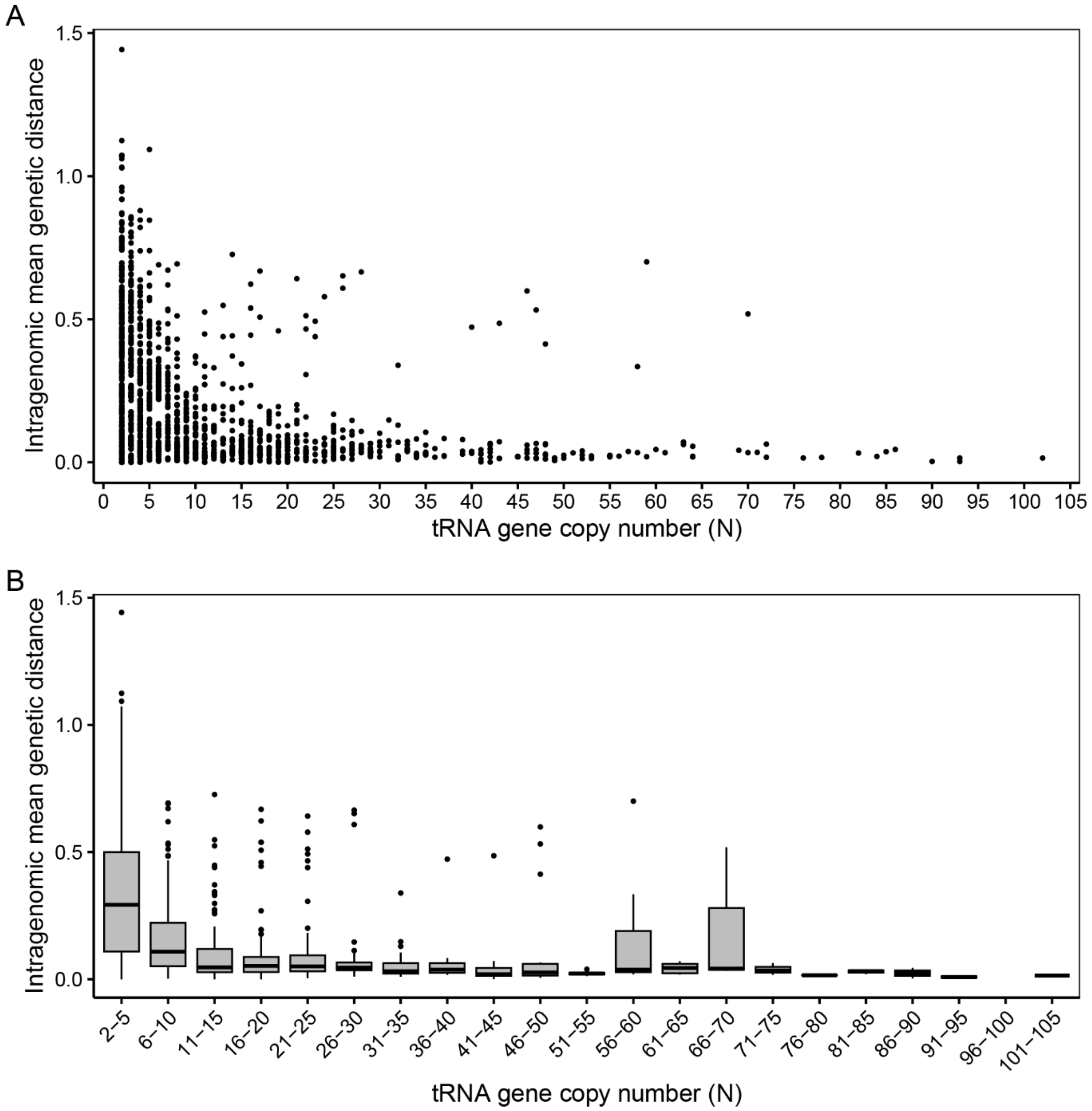


**Figure S7.** Intragenomic sequence polymorphism of tRNA genes in relation to copy-number (A) and by categorical groups (B). The genetic distance was calculated for each tRNA gene detected in a single genome. Each data point corresponds to the sequence variability among all copies for the same tRNA gene present in each genome.

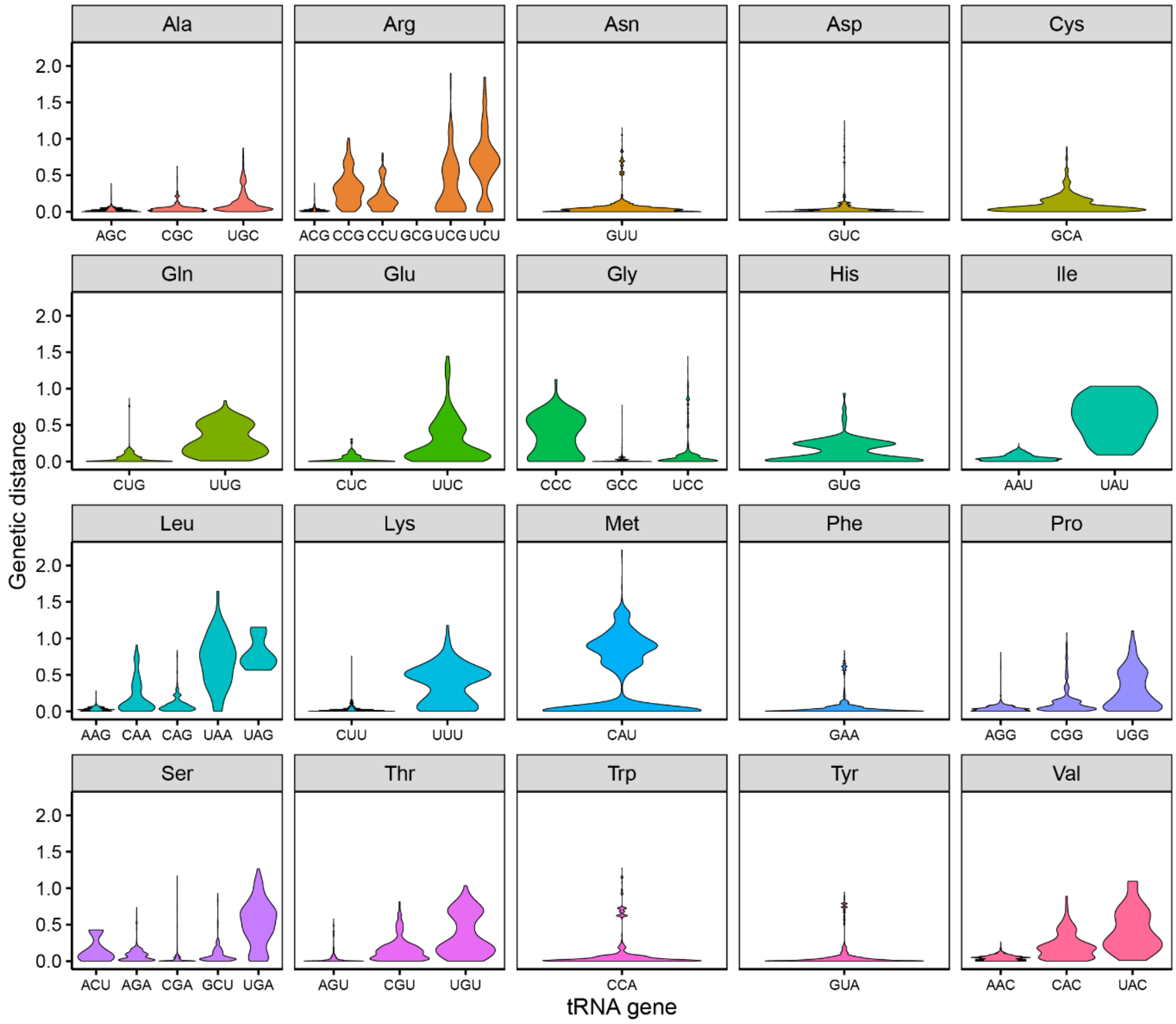


**Figure S8.** Intragenomic sequence variation of tRNA genes by anticodon type and respective amino acid.

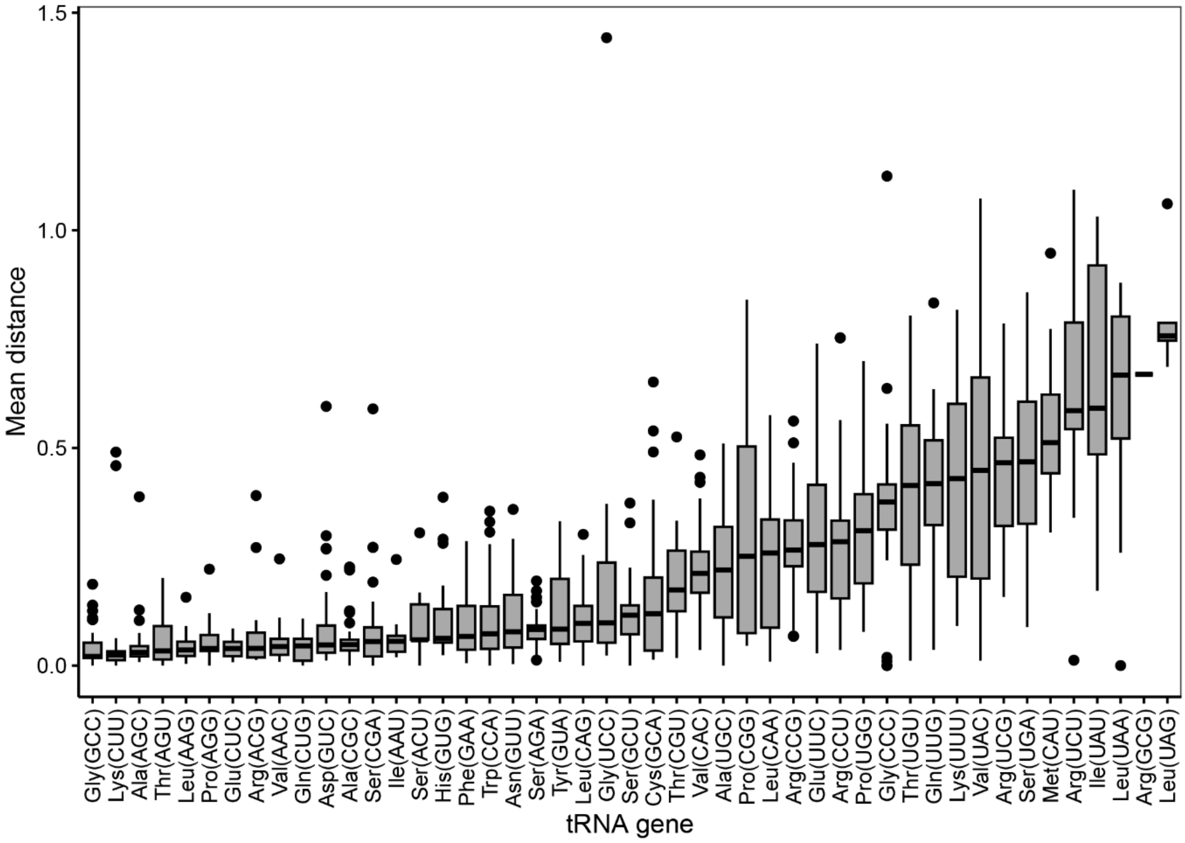


**Figure S9.** Mean genetic distance among tRNA genes by anticodon and respective amino acid across all Trichosporonales species. The plot is sorted by median value.

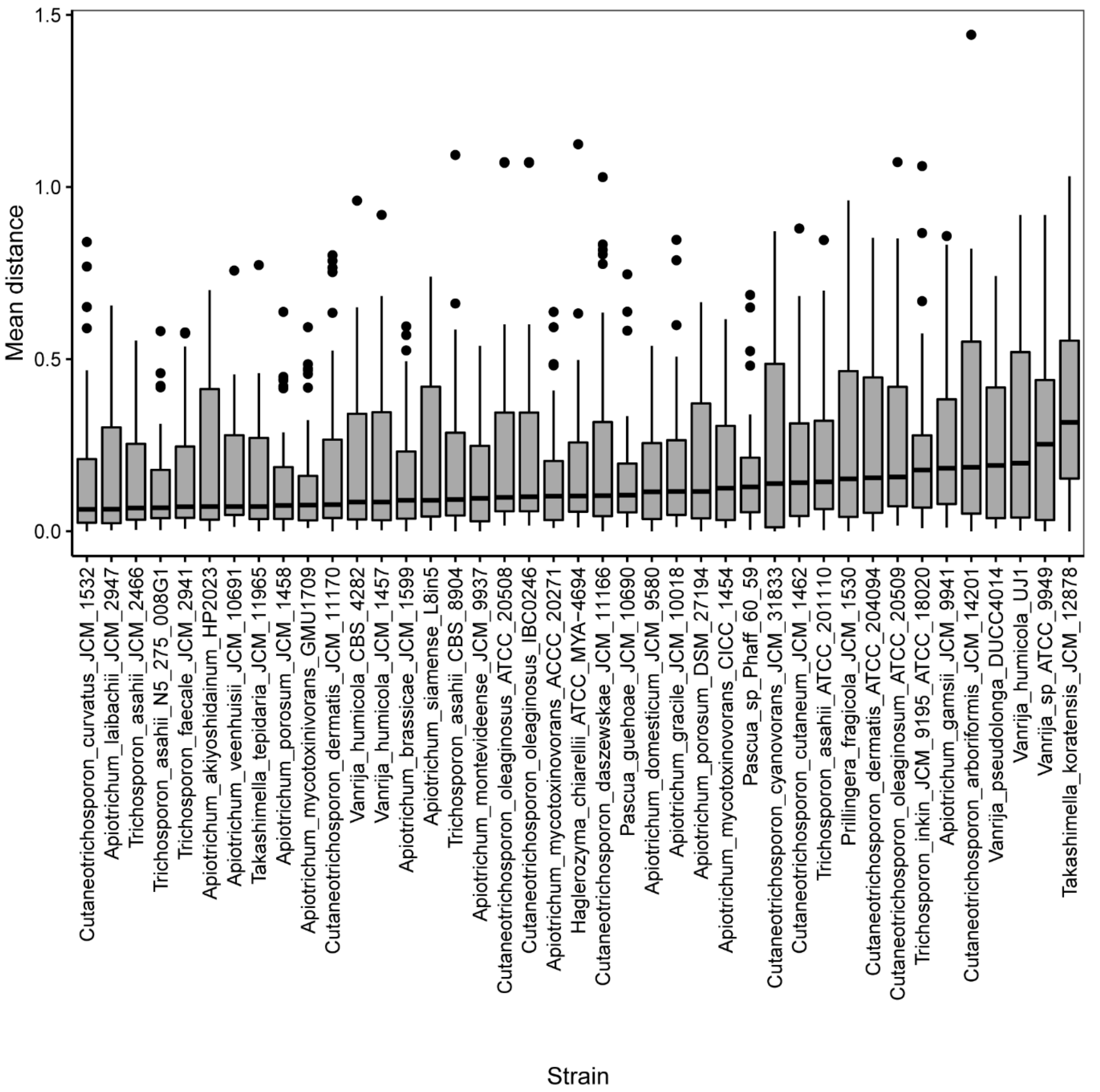


**Figure S10.** Mean genetic distance among tRNA genes by genome assemblies. The plot is sorted by median value.

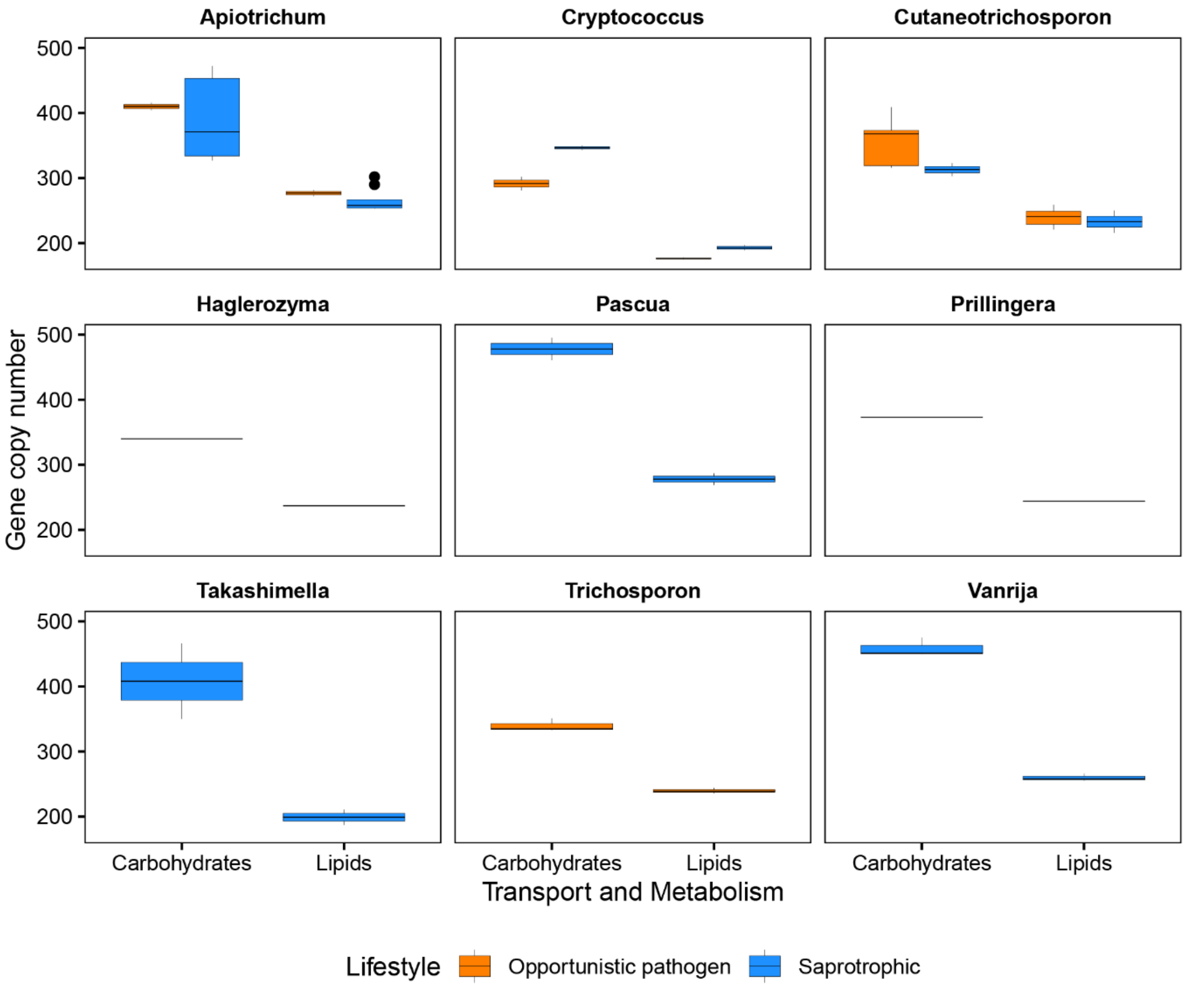


**Figure S11.** Gene copy number for genes involved in lipid and carbohydrate transport and metabolism. Gene copy numbers are compared between genera and opportunistic pathogenic and saprotrophic lifestyles.

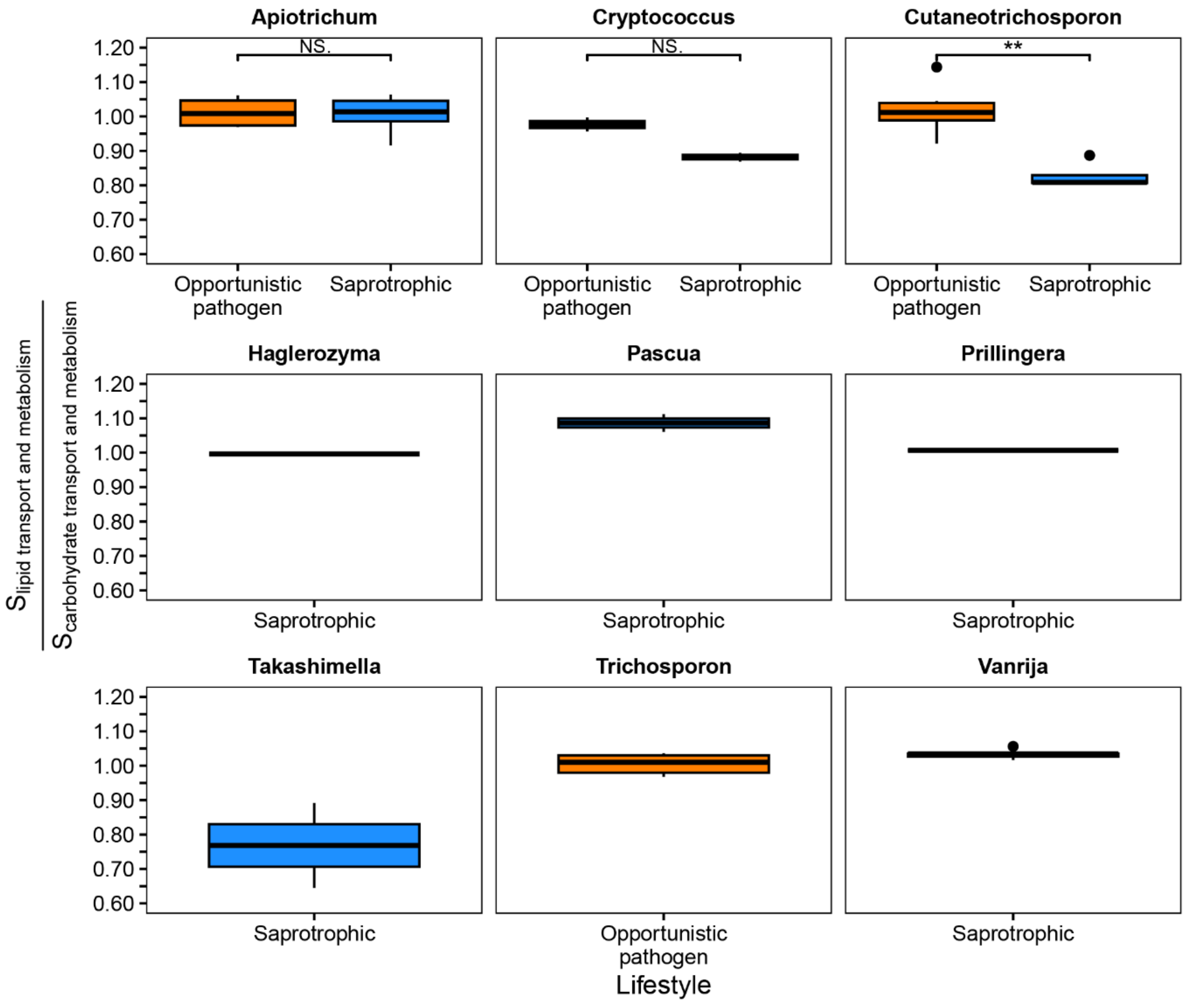


**Figure S12.** Relative translation efficiency for genes involved in lipid and carbohydrate transport and metabolism. Relative translation efficiency (S index) is compared between genera and opportunistic pathogenic and saprotrophic lifestyles.

25. Bive BZ, Sacheli R, Situakibanza Nani-Tuma H, Kabututu Zakayi P, Ka A, Mbula Mambimbi M, et al. Clinical epidemiology and high genetic diversity amongst *Cryptococcus* spp. isolates infecting

people living with HIV in Kinshasa, Democratic Republic of Congo: Von Ewigkeit zu Ewigkeit. PLoS One. 2022;17:e0267842. doi:10.1371/journal.pone.0267842.

33. Wang L, Guo S, Zeng B, Wang S, Chen Y, Cheng S, et al. Draft Genome Assembly and Annotation for Cutaneotrichosporon dermatis NICC30027, an Oleaginous Yeast Capable of Simultaneous

Glucose and Xylose Assimilation. Mycobiology. 2022;50:69–81. doi:10.1080/12298093.2022.2038844.
